# Distribution of prophage-encoded Pas sRNAs across pathogenic *Escherichia coli*

**DOI:** 10.64898/2026.05.13.724894

**Authors:** Dennis X. Zhu, Svetlana A. Shabalina, Gisela Storz

## Abstract

Numerous base pairing small RNAs (sRNAs), which are an integral part of regulatory networks in bacteria, are encoded on mobile genetic elements (MGEs) in pathogenic strains of *Escherichia coli*. These sRNAs help coordinate the expression of MGE-encoded virulence factors with core genome-encoded cellular pathways. To investigate the evolution of MGE-encoded sRNAs, we queried public databases to characterize the distribution of PasA, PasB, PasC, PasD1, and PasD2, five prophage-encoded sRNAs discovered in enteropathogenic *E. coli*. We find that while the Pas sRNAs are largely restricted to pathogenic lineages of *Escherichia* and *Shigella*, they exhibit diversity in sequence, genomic presence, and copy number across strains. Based on phylogenetic analysis, the Pas sRNAs originate from multiple ancestral lineages and associate with specific *E. coli* pathovars, consistent with horizontal acquisition followed by retention. Syntenic analysis suggests a phage origin for the Pas sRNAs, likely from Shiga-toxin encoding phages, but the sRNAs appear to have diverged substantially following their integration into bacterial chromosomes. Comparative and structural analyses further suggest that the PasA and PasC sRNAs share a common ancestor as is the case for PasD and STnc100, another prophage-encoded sRNA. These findings add to our understanding of how accessory genome-encoded sRNAs emerge and evolve.

## Introduction

Bacterial small RNAs (sRNAs) are a diverse group of regulatory RNAs that are widespread in bacteria and predominantly regulate gene expression post-transcriptionally by base-pairing with messenger RNA (mRNA) targets (reviewed in [1,2]). The most well-studied class of base-pairing sRNAs regulate trans-encoded mRNA targets through limited sequence complementarity with their targets. In many organisms, these sRNAs require the activity of an RNA-binding protein chaperone such as Hfq or ProQ to facilitate base-pairing but are able to interact with multiple, different mRNA targets, forming interconnected regulatory networks (reviewed in [3]). sRNA regulation has been implicated in numerous critical cellular processes in bacteria including the regulation of central carbon metabolism, iron homeostasis, the cell envelope, quorum sensing, and virulence (reviewed in [4]).

Despite their broad importance for bacterial physiology, relatively little is known about the evolution of the sRNA regulators. The distribution and evolution of sRNAs across bacteria has historically been difficult to study due to their short sequences and lack of conservation across broad evolutionary distances (reviewed in [5]). The relative lack of sequence constraints facilitates the *de novo* emergence of sRNAs from diverse genetic contexts [6] but also allows them to be rapidly lost from bacterial genomes [7]. Most non-coding sRNA families are conserved only to the genus or family level, whereas protein-coding RNAs are often conserved across the bacterial phylum [8]. Evolutionary studies also are limited by the pool of documented sRNAs from only a subset of bacterial genomes. Previous studies examining sRNA evolution in enteric bacteria relied on sRNA lists compiled from the reference strains *Escherichia coli* K-12 and *Salmonella enterica* serovar Typhimurium ST4/74 and showed that, with the exception of several highly conserved sRNAs such as RyhB, few *E. coli* sRNAs are conserved beyond the Enterobacterales [7,9,10]. Recent studies in *Vibrio* and *Pseudomonas* also documented that many sRNAs are genus-specific but show diversity across strains and species [11,12]. Interestingly, different lineages of bacteria regulate the same physiological process even though the sequences can be very different, as was discovered for the sRNA regulators of iron homeostasis in *Escherichia* and *Pseudomonas* (reviewed in [13]). Given this limited conservation, comparing sRNAs within narrower phylogentic windows such as at the strain- or species-level may offer more insight into their evolution.

The application of high-throughput sequencing approaches has facilitated the discovery of novel sRNAs [14,15] (reviewed in [16]). As the number of known bacterial sRNAs has increased, so too has our understanding of the mechanisms underlying their evolution (reviewed in [17]). Horizontal gene transfer is a major driver of bacterial genome evolution and has been shown to contribute to both the *de novo* emergence of novel sRNAs [18] and the introduction of existing sRNAs into genomes (reviewed in [19]). The *E. coli* K-12 genome, which is 4.6 Mb and encodes roughly 4,288 genes, represents only a fraction of the genetic pool within the *E. coli* pangenome, which is estimated to include as many as 75,000 genes (reviewed in [20]). *E. coli* genomes share a conserved “core genome” comprised of roughly 2,200 genes, while mobile genetic elements (MGEs) including plasmids, bacteriophages, and chromosomal insertion sequences comprise the “accessory genome” that is distributed across strains. MGEs are a driver in the evolution of both pathogenesis and sRNAs in *E. coli* (reviewed in [19]). Pathogenic strains of *E. coli* are thought to have emerged through stepwise acquisitions of virulence factors on MGEs that have stably integrated into the genome (reviewed in [20]). These virulence factor-encoding MGEs have served as both a regulatory target of core genome-encoded sRNAs and as a reservoir of accessory genome sRNAs.

Unlike core genome sRNAs which are encoded in essentially all *E. coli* genomes, accessory genome sRNAs show variable distribution across strains. As a result, sRNA-based post-transcriptional regulation can vary greatly between different strains of the same bacterial species depending on the repertoire of sRNA genes that are present in the genome [12]. This point is illustrated by the observation that, in the enterohemorrhagic *E. coli* (EHEC) strains Sakai and EDL933, deletion of *hfq* results in increased expression of virulence genes from the locus of enterocyte effacement (LEE) pathogenicity island [21]; while deletion of *hfq* in another EHEC strain 86-24 had the opposite effect, repressing expression of LEE genes [22]. In addition to regulating MGE genes, many MGE-encoded sRNAs regulate the expression of core genome genes. As an example, the EHEC-specific prophage-encoded sRNA Esr41 represses the expression of LEE virulence genes while simultaneously activating motility through induction of core genome encoded FliC (flagellin) expression [23]. These examples highlight the value of understanding the distribution of accessory genome sRNAs across different bacterial strains.

To explore the principles underlying the evolution of sRNA genes in pathogenic *E. coli*, we examined the distribution of five prophage-encoded sRNAs genes recently detected in the enteropathogenic *E. coli* (EPEC) strain E2348/69 [24]. These novel sRNAs were identified using RIL-seq (RNA interaction by ligation and sequencing), an approach that identifies interacting sRNA-mRNA pairs by ligating and then sequencing RNA species bound to Hfq [15,25]. The five sRNAs, which were among the most abundantly detected sRNAs in E2348/69 and were validated by northern blot analysis, are named PasA, PasB, PasC, PasD1, and PasD2 (Pas for <u>P</u>athogenesis <u>a</u>ssociated <u>s</u>RNA for their location within prophage regions). The Pas sRNA genes are all located intergenically within lambdoid prophage regions on the E2348/69 chromosome [26]. Despite their common name, the Pas sRNAs were originally described as distinct genes, though near-complete sequence identity between PasD1 and PasD2 and partial sequence similarity between PasA and PasC was noted. The function of these sRNAs has not been elucidated, but RIL-seq analysis indicates that they interact with numerous accessory and core genome-encoded mRNAs.

Here, we searched for Pas sRNA genes in public databases of sequenced bacterial genomes to characterize their distribution and found that the Pas sRNA sequences are largely restricted to *Escherichia* and *Shigella* species and enriched in pathogenic strains of *E. coli*. We further mapped the presence or absence of Pas sRNAs to a strain-level phylogeny and performed synteny analysis. These comparisons led us to conclude that the Pas sRNAs were likely carried into certain pathogenic lineages via Shiga toxin-containing bacteriophages. Based on the similarity of their sequences and neighboring gene contexts, we conclude that PasA/PasC are related gene variants as are PasD1/PasD2 together with STnc100, another prophage-encoded sRNA. Additionally, PasA/PasC and PasD1/PasD2 appear to have been repeatedly acquired and maintained in several EPEC and EHEC lineages. In contrast, PasB is distributed more broadly and is frequently found in extraintestinal pathogen isolates. This study helps expand our understanding of how prophage-encoded accessory sRNAs are distributed across pathogenic *E. coli* strains.

## Materials and methods

### BLAST searches

Nucleotide Basic Local Alignment Search Tool (BLAST) searches for Pas sRNA genes were performed using BLAST+ v.2.6.0 ‘blastn’ with DNA sequences of the five Pas sRNA genes (PasA, PasB, PasC, PasD1, and PasD2) from the *Escherichia coli* O127:H6 strain E2348/69 (GCF_00026545.1) [26] as the query. Default parameters were used for all searches unless otherwise stated. The search for Pas sRNA genes in bacterial genomes was performed against the NCBI ‘prok_complete_genomes’ database on April 18, 2025 using a word size of 28. To identify more divergent hits, additional blastn searches were performed with relaxed parameters (a word size of seven and automatically adjusted parameters for short sequences) against the same database while excluding sequences from *Escherichia* and *Shigella* taxonomic groups. The search for Pas sRNA genes in bacteriophage genomes was performed against the NCBI ‘Complete_Bacteriophages’ database on May 05, 2025 and against the Metagenomic Gut Virus catalogue [27] using a word size of 11. Bacterial genome results were filtered post-search for hits with a ‘qcovs’ value greater than 75, and bacteriophage genome results were filtered for hits with a ‘qcovs’ value greater than 50.

Genome sequence and annotation data for the 1,362 bacterial genomes containing at least one Pas sRNA gene hit was downloaded using NCBI Datasets [28]. BLAST+ v2.6.0 was used to create a local database (‘pas_1362’) from the 1,362 genomic DNA sequences. The ‘pas_1362’ BLAST database was used as the subject for blastn searches within the ‘pas_1362’ genome set.

### RNA structure predictions and analysis

RNA secondary structures were predicted from nucleotide sequences using the RNAfold program from the ViennaRNA Package v.2.0 [29]. Multiple sequence alignments (MSA) conduscted by MUSCLE v.5 [30] were folded using RNAalifold (from ViennaRNA 2.7.0) [29]. Afold and Hybrid [31,32] were used to predict locally folded secondary structures and potential RNA-RNA duplex elements within clusters/MSAs of sRNA candidates. All structure predictions were performed using default parameters, and the minimum-free energy prediction is shown. Infernal v.1.1.5 [33] was used to compare candidate RNA structures against Rfam covariance models (CM), build and calibrate new covariance models (*cmscan*, *cmbuild*, *cmcalibrate*) for separate clusters of Pas RNAs, and perform structure-informed homology searches (*cmsearch*). Known sequence profiles and CMs were downloaded from the Rfam database [34] on March 2024. Comparative analysis and MSAs of isolated RNA candidates were conducted using *cmalign* and MUSCLE v.5 [30], with pairwise comparisons further refined using the OWEN program [35,36].

Phylogenetic relationships were inferred using the Maximum Likelihood method under multiple substitution models in MEGA X [37,38]. The evolutionary history was inferred by using the Maximum Likelihood method and Kimura 2-parameter model [39]. Initial trees for the heuristic search were obtained automatically by applying Neighbor-Join and BioNJ algorithms to a matrix of pairwise distances estimated using the Maximum Composite Likelihood (MCL) approach, and then selecting the topology with superior log likelihood value. A discrete Gamma distribution was used to model evolutionary rate differences among sites (see Figure legends for details).

### Strain classification and MLST typing

Bacterial genomes in the ‘pas_1362’ dataset were classified to the genus and species level using associated GenBank taxonomic information. Strain- and serotype-level classification was performed by filtering for specific NCBI taxonomy id numbers (txid): ‘txid83334’ or ‘txid1045010’ for ‘*E. coli* O157’ and ‘txid1055538’ or ‘txid1078034’ for ‘*E. coli* O145’. The total number of sequenced genomes belonging to each taxon was characterized using metadata downloaded using NCBI datasets filtered for specific txid numbers [40].

Strains were classified into pathovars based on the presence of specific virulence factor genes in the genome: enteropathogenic *E. coli* (EPEC) was defined by *eae* (Gene ID: 915471); Shiga-toxin producing *E. coli* (STEC) was defined by *stx1A* (Gene ID: 916678) or *stx2A* (Gene ID: 912807); enterohemorrhagic *E. coli* (EHEC) was defined by the presence of both *eae* and *stx1A* or *stx2A*; enteroaggregative *E. coli* (EAEC) was defined by *aggR* (Gene ID: 7169332); Stx-EAEC was defined by the presence of both *aggR* and *stx1A* or *stx2A*; and enterotoxogenic *E. coli* (ETEC) was defined by *eltA* (Gene ID: 8571459) or *eltB* (Gene ID: 8571464). Isolates lacking any of the listed virulence factor genes were classified as ‘Other’. The presence of virulence factor genes was detected by blastn search against the ‘pas_1362’ BLAST database.

Multi-locus sequence typing was performed using the seven gene Achtman scheme [41]. DNA sequences of seven housekeeping genes (*adk*, *fumC*, *gyrB*, *icd*, *mdh*, *purA*, *recA*) were extracted from each genome in the ‘pas_1362’ dataset based on feature boundaries defined by the NCBI Prokaryotic Genome Annotation Pipeline (PGAP) [42]. Genomes missing or encoding more than one copy of any of the seven MLST genes were excluded from the analysis, leaving a total of 1,211 genomes with unambiguous MLST sequences. MLST gene sequences were uploaded to PubMLST to determine the sequence type of each bacterial isolate [43].

### Maximum-likelihood tree construction

Maximum-likelihood phylogenetic trees were constructed using the concatenated DNA sequence of the seven housekeeping MLST genes. MLST sequences were aligned with Muscle v.5 using the Super5 algorithm [30]. Phylogenetic relationships between bacterial isolates were inferred by maximum-likelihood estimation using RAxML (Randomized Axelerated Maximum Likelhood) v.8 using the GTRCAT approximation and 200 bootstraps [44]. The final tree was re-rooted at the node separating the divergence of *Escherichia albertii* isolates from the rest of the branches. Phylogenetic trees were visualized using the ‘ggtree’ R package [45].

### Clustering and comparison of flanking regions

Custom R code was used to extract the DNA sequences of roughly 20 kbp regions consisting of 10 kbp flanks surrounding each Pas sRNA gene hit in the ‘pas_1362’ dataset. Flanking sequences were aligned and clustered using MMseqs2 using ‘-cov-mode 0-cluster-mode 0’ with a mimium coverage of 75% and minimum sequence identity of 20% [46]. Flanking region synteny was visualized using the ‘geneviewer’ R package v.0.1.10 (https://nvelden.github.io/geneviewer/authors.html#citation). Features were compared across regions based on manual inspection of PGAP annotations.

### Prophage prediction

PHASTER (PHAge Search Tool – Enhanced Release) was used to identify prophage regions within bacterial genomes [47]. The best matching phage was determined based on blastp search of protein-coding genes within the prophage region and filtered for phages with a minimum of 20% similar proteins. Prophage completeness was classified based on default PHASTER settings.

## Results

### Pas sRNAs show sequence diversity across bacteria

To characterize the distribution of Pas sRNAs across all bacteria, we performed a blastn search of the five Pas genes (PasA, PasB, PasC, PasD1, and PasD2) found in *E. coli* (EPEC) strain E2348/69 [24] against the database of 147,705 complete prokaryotic genomes available on NCBI (‘prok_complete_genomes’ as of April 18, 2025) (**Fig. 1A**). The BLAST search identified 4,600 hits across 1,362 unique bacterial genomes (**Supplementary Table S1A**) (‘pas_1362’). Nearly all of the hits (4,588/4,600) mapped to bacterial chromosomes, though 12 mapped to plasmids (**Supplementary Table S1B**). Plasmids in the dataset typically encode a single copy of a Pas sRNA gene, except one plasmid that encodes a copy of both PasA and PasD1.

**Figure 1.**
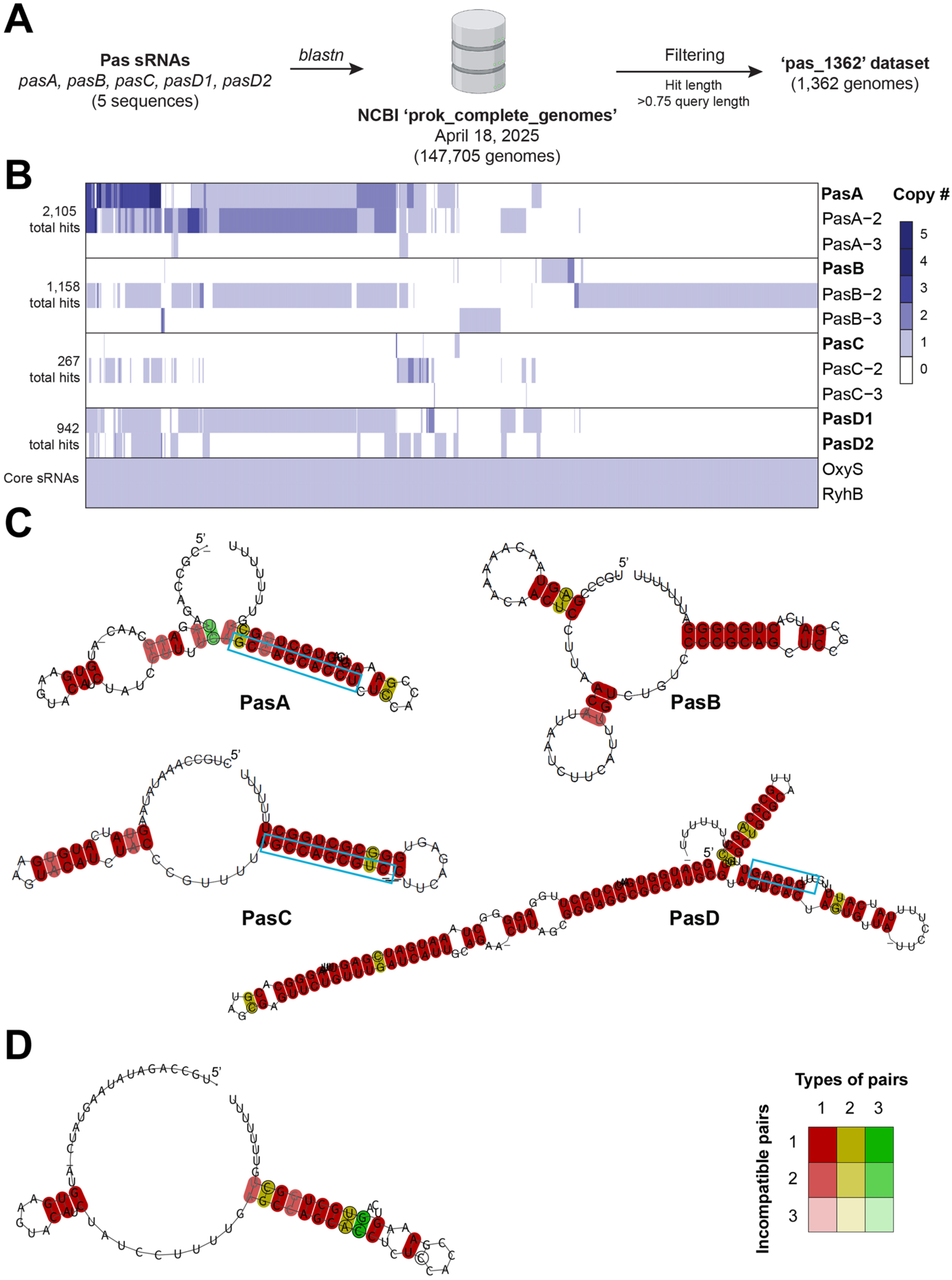
Bioinformatic analysis reveals sequence variation in Pas sRNA genes across bacteria. (A) Overview of bioinformatic approach. All five Pas sRNA nucleotide sequences from *E. coli* E2348/69 were searched against the NCBI complete prokaryotic genomes database (‘prok_complete_genome’) using *blastn*. The search identified 1,362 unique bacterial genomes encoding at least one Pas sRNA. These genomes were stored as a stable database (‘pas_1362’) for downstream analyses. (**B**) Heatmap showing the gene copy number of each Pas sRNA gene or sequence variant, as well as the core OxyS and RyhB sRNA genes, in the ‘pas_1362’ dataset. Each column represents a single bacterial genome, and each row represents a different gene or variant. Only the most prevalent variants for each gene are shown. Columns are clustered based on sRNA profile similarity. The total number of BLAST search hits for PasA, PasB, PasC, and PasD is shown. (**C**) Predicted consensus secondary structures based on MSAs of PasA, PasB, PasC, and PasD. (**D**) Predicted consensus secondary structure based on MSAs for PasA and PasC together. For both (C) and (D), RNAalifold [29] was applied to the Pas MSAs. Colors are based on the MSAs of each Pas RNAs (**Supplementary Fig. S1**), with darkest colors indicating all RNAs in the alignment have the predicted base pair, though this can be through one (red), two (yellow) or three (green) different combinations. If a predicted base-pair is formed by several different nucleotide combinations, then coordinated or compensatory mutations have taken place. Pale colors indicate that a base-pair cannot be formed in indicated number sequences of the alignment. Nucleotides predicted to be involved in intermolecular interactions (seed regions) are shown in blue boxes.

In contrast to core genome-encoded sRNAs, which are present uniformly across isolates of a given species, the Pas sRNAs are distributed unequally across bacterial genomes in the ‘pas_1362’ dataset (**Fig. 1B**). Through our BLAST search, we were able to identify several sequence variants of the Pas sRNA genes that we queried (**Fig. 1C, Supplementary Fig. S1** and **S2**). Each Pas gene had multiple variants that represented >1% of the total hits, suggesting that these variants have become evolutionarily fixed across the population. In fact, for PasA, PasB, and PasC, the query gene sequence from *E. coli* strain E2348/69 was not the most prevalent sequence variant across the dataset. For the sake of clarity, Pas sRNAs that were previously characterized in E2348/69 are referred to by their original names (i.e. PasA, PasB, PasC, PasD1, and PasD2) while other sequence variants discussed here are named with dash numbers in order of their prevalence in the dataset (e.g. PasA-2, PasA-3). PasD1 and PasD2 differ by four nucleotides, so we considered them sequence variants of the same sRNA and thus for some analyses will refer to them collectively as ‘PasD’.

PasA variants are most prevalent in the dataset (2,105 total hits) and often are found as multiple copies in a single genome. PasB variants are the second most prevalent (1,158 total hits) followed by PasD variants (942 total hits) and PasC variants (267 total hits). PasB is frequently found alone as the only Pas sRNA gene in a genome. In contrast, 96.2% of genomes encoding at least one copy of PasA also encoded at least one copy of PasD (Phi coefficient = 0.92) (**Supplementary Table S2A)**. Typically, only a single variant of PasB or PasC is present within a given bacterial genome. However, numerous genomes encode multiple different variants of PasA and PasD, which may possess non-redundant regulatory functions. Collectively, the heterogeneous distribution of Pas sRNA genes and sequence variants across bacterial isolates generates diversity in sRNA repertoire.

Unlike protein-coding RNAs, whose evolution is largely constrained by codon requirements, sRNA sequence evolution is primarily shaped by intra- and intermolecular interactions. These constraints are critical for maintaining RNA secondary structure, protein binding sites, and conserved regions, often called seed sequences, that nucleate base-pairing with target RNAs. To examine how the sequence polymorphisms in Pas sRNA variants affect RNA structure, we aligned the variant sequences for each Pas sRNA and used *RNAalifold* to predict their consensus secondary structure (**Fig. 1C and Supplementary Fig. S1**) [48]. A core consensus RNA fold was detected for each Pas sRNA dataset. All the Pas sRNAs show a 3’ stem-loop followed by a poly-uridine tract that is indicative of an intrinsic transcription terminator and likely an Hfq-binding site.

PasA sequence variants display significant differences in sequence. PasA and PasA-2, the most common variants distributed almost evenly across our datasets, share a conserved 3’ end but diverge in the first 40 bp of the gene (**Supplementary Fig. S1A**). Previous northern blot analysis of PasA suggested that the 5’ end of the mature 84 nt PasA sRNA is generated by cleavage from a longer transcript [24]. Due to potential differences in processing, we cannot confidently predict the position of the PasA-2 5’ end. Thus, the structure of PasA-2 was predicted using an 84 nt sequence to match the size of the processed PasA-2 sRNA. Although PasA and PasA-2 diverge most in nucleotide sequence, there is a conserved RNA folding core (**Fig. 1C**) with some suboptimal variations at the 5’ ends (**Supplementary Fig. S2A**). PasA-3 is identical in sequence to PasA, except that it is lacking a 3’ intrinsic terminator structure. No terminator sequence could be identified within 100 bp downstream of the PasA-3 sequences, so PasA-3 may be a pseudogene or an ancestor sequence for the functional PasA sRNA. Overall, the sequence differences across PasA sRNA variants are likely to affect the levels, processing, structure, and function of this sRNA family.

Interestingly, PasA variants can be unambiguously aligned with PasC variants, and a consensus RNA fold can be predicted with the MSA of all PasA and PasC sRNAs variants (**Fig. 1D** and **Supplementary Fig. S1B**). The similarity between PasA and PasC at the sequence and structural levels indicate a common origin of these sRNAs.

In contrast to PasA, other Pas sRNAs only show a few nucleotide differences. However, even minor sequence variations are predicted to lead to some differences in optimal and suboptimal RNA structures. For example, a single nucleotide change between PasB and PasB-2 at position 27 (**Supplementary Fig. S2B**), can remove the predicted consensus stem-loop in the optimal PasB structure. Similarly, single nucleotide polymorphisms may alter the position or number of predicted 5’ stem-loop(s) in the optimal folding of PasC-2 and PasC-3, respectively, as compared to PasC (**Supplementary Fig. S2C**). Sequence polymorphisms between PasD1 and PasD2 have little effect on their predicted structure. However, RIL-seq analysis indicates that PasD1 and PasD2 bind overlapping but distinct sets of targets [24] demonstrating that small sRNA sequence variations in the seed region can potentially alter regulatory specificity without large changes in sequence or folding (**Supplementary Fig. S2D**).

Pearl Mizrahi *et al.* predicted the seed sequence positions for PasA and PasD based on conserved, complementary sequences in their RIL-seq targets [24]. These predicted seed sequences are conserved across PasA and PasD variants in our dataset (**Fig. 1C** and **Supplementary Fig. S1A**), though the observation that the PasA seed overlaps the terminator stem is unusual. We next compared the seed region of the PasA with the corresponding conserved region in the PasC variants (**Supplementary Fig. S1A**). Given some differences between the sequences, we suggest that despite their common origin, PasA and PasC sRNAs may interact with distinct sets of targets. Based on the most conserved sequences in the MSA, we propose the PasB seed is in a canonical single-stranded region just upstream of the terminator stem loop sequence.

### Pas sRNAs are overrepresented in EPEC and EHEC strains

Pathogenicity in *E. coli* has evolved heterogeneously across different phylogenetic lineages, likely due to differences in the ability of certain *E. coli* strains to acquire and maintain virulence factors encoded on MGEs. Since we observed that Pas sRNAs also were heterogeneously distributed across bacterial isolates, we asked whether certain bacterial species or strains were overrepresented in the ‘pas_1363’ dataset. To this end, we classified the ‘pas_1362’ genomes into species based on their NCBI annotations. All of the bacterial genomes in the ‘pas_1362’ dataset belong to either the genus *Escherichia* or *Shigella*, with a majority (95.4%; 1,299/1,362) of the hits classified as *E. coli* (**Fig. 2A**). *Escherichia albertii* is the most distantly related taxon that encodes a Pas sRNA. Pas sRNA genes show heterogeneous species- and lineage-specific prevalence. PasA is the only Pas sRNA found in *E. albertii* while PasB is more prevalent within *Shigella* genomes than the other Pas sRNAs. Overall, Pas sRNAs appear largely restricted to *Escherichia* and *Shigella* clades as relaxed similarity searches for more divergent homologs in other bacterial lineages did not yield significant matches (see Materials and Methods).

**Figure 2.**
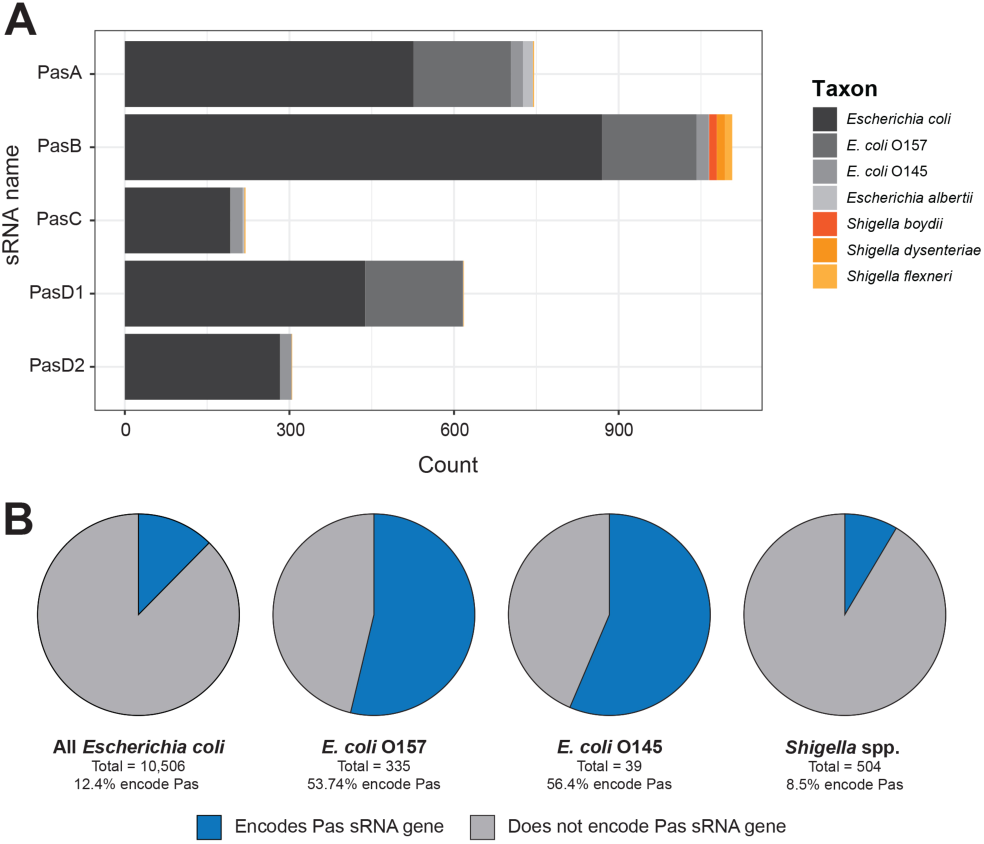
Pas sRNAs are primarily found within pathogenic *E. coli* isolates. (**A**) Bar graph showing the species distribution of bacterial genomes containing each Pas sRNA. Genomes are classified by their NCBI species annotation. *E. coli* isolates are further classified into O157 and O145 serotypes. All other serotypes and non-serotyped isolates are included in ‘*Escherichia coli*’. (**B**) Pie graphs showing the percentage of NCBI genomes (available as of April 18, 2025) that encode at least one Pas sRNA for the taxa: ‘All *Escherichia coli*’ (txid562), ‘*E. coli* O157’ (txid83334 and txid1045010), ‘*E. coli* O145’ (txid1055538 and txid1078034), and ‘*Shigella* spp.’ (txid620).

To obtain a more granular view of Pas sRNA gene distribution in *E. coli*, we specifically examined *E. coli* genomes belonging to the O157 and O145 serotypes, which are the two most prevalent *E. coli* serotypes in the ‘pas_1362’ dataset and are both notable for causing severe enterohemorrhagic disease in humans. Among sequenced genomes available on NCBI, Pas sRNAs are overrepresented (hypergeometric *p* < 0.05) in *E. coli* O157 (50.7%) and *E. coli* O145 (56.4%) isolates compared to *E. coli* genomes as a whole (12.4%). PasC and PasD2 are completely absent from *E. coli* O157 isolates while PasD1 is absent from *E. coli* O145 isolates (**Fig. 2A**), indicating that these two EHEC lineages evolved with different repertoires of accessory genome-encoded Pas sRNAs.

The association between Pas sRNAs and certain pathogenic serovars of *E. coli* led us to examine the distribution of Pas genes within different *E. coli* pathovars. We classified the 1,299 *E. coli* isolates in the ‘pas_1362’ dataset into pathovar groups based on the presence of definitive virulence genes (*eae* for enteropathogenic *E. coli* [EPEC]; *stx1A*/*stx2A* for Shiga toxin producing *E. coli* [STEC]; both *eae* and *stx1A*/*stx2A* for enterohemorrhagic *E. coli* (EHEC); *aggR* for enteroaggregative *E. coli* [EAEC]; *aggR* and *stx1A*/*stx2A* for EAEC-Stx; and *eltA*/*eltB* for enterotoxigenic *E. coli* [ETEC]). Isolates lacking these specific virulence factors were classified as ‘Other’, although they still may be pathogenic. The pathovar identity of a strain was significantly associated with certain repertoires of Pas sRNA genes (Fisher’s exact test p < 0.0005). Specifically, EPEC and EHEC genomes were most likely to encode multiple copies of PasA and one copy of both PasB and PasD (**Supplementary Table S2C).** PasA, PasC, and PasD variants were found almost exclusively within EPEC and EHEC isolates (**Supplementary Fig. S3A**) defined by the locus of enterocyte effacement (LEE) pathogenicity island, though none of the Pas sRNA genes are directly encoded within the LEE island as indicated by the distance to the LEE-encoded *eae* and *stx* genes (**Supplementary Fig. S3B**). Thus, the Pas sRNAs likely were not acquired concomitantly with the LEE but are preferentially acquired or maintained in EPEC and EHEC strains through other mechanisms. In contrast, strains classified as ETEC, EAEC, or ‘Other’ most commonly encoded just one copy of PasB as the only Pas sRNA gene (**Supplementary Table S2C)**. These data suggest that PasA, PasC and PasD are genetically linked while PasB entered *E. coli* genomes independently.

### Phylogenetic analysis supports horizontal evolution of Pas sRNA genes

We next examined the phylogenetic distribution of Pas genes. We constructed a maximum-likelihood tree of the bacterial genomes in the ‘pas_1362’ dataset using seven housekeeping genes from the Achtman Multilocus Sequence Typing (MLST) scheme (*adk*, *fumC*, *gyrB*, *icd*, *mdh*, *purA*, and *recA*) [41]. MLST genes could be unambiguously identified on the chromosome of 1,211 genomes. We constructed a maximum-likelihood tree using these 1,211 MLST sequences (**Supplementary Fig. S4**) and a reduced tree using 271 non-identical MLST sequences, with 970 identical sequences removed, to reflect the phylogenetic relationships of the genomes in the ‘pas_1362’ dataset (**Fig. 3**). *E. coli* strains are assigned into clonal complexes by PubMLST based on sequence similarity of MLST genes to a central allelic sequence type profile [43]. The two most highly represented clonal complexes in the ‘pas_1362’ dataset are ST11 (n = 347), which includes serotype O157:H7 EHEC strains, and ST131 (n = 304), a multi-drug resistant *E. coli* lineage [49], which are the two strains most frequently associated with intestinal and extraintestinal *E. coli* infection, respectively.

**Figure 3.**
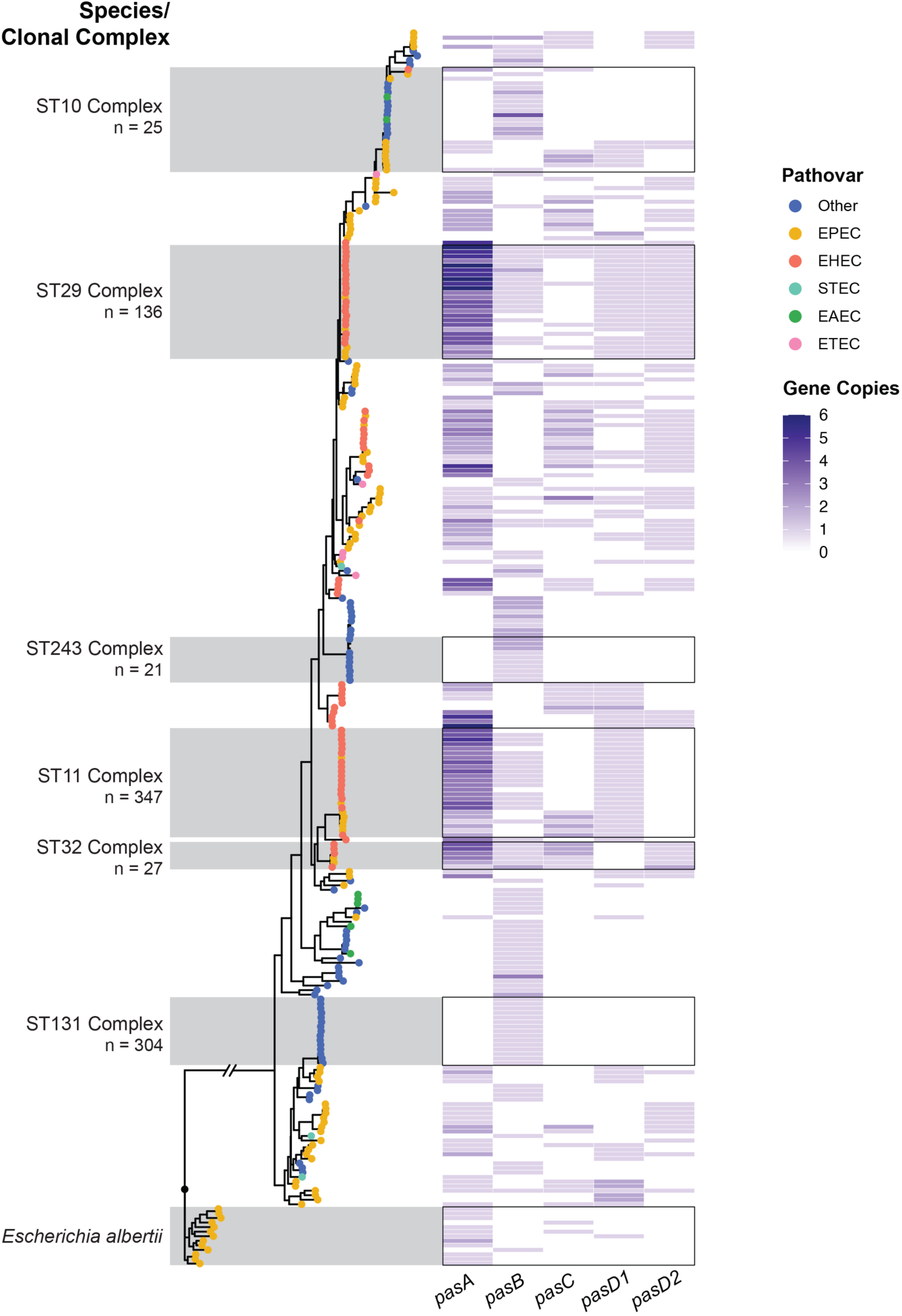
Maximum likelihood tree of bacteria isolates from the ‘pas_1362’ dataset. Phylogenetic estimations were constructed from the nucleotide sequence of seven housekeeping genes from the Achtman Multilocus Sequence Typing (MLST) scheme (*adk*, *fumC*, *gyrB*, *icd*, *mdh*, *purA*, *recA*). RAxML was used for maximum likelihood estimation using the ‘GTRCAT’ substitution model. MLST sequences could be identified for 1,211 out of 1,319 *Escherichia* genomes in the dataset. The final tree displays 271 genomes with non-identical MLST sequences (940 genomes had identical MLST sequences and were trimmed). The tree is rooted at the divergence of *Escherichia albertii* from *E. coli* isolates. Branch tips are colored based on the pathovar classification of each bacterial isolate (see Materials and Methods). Briefly, *E. coli* genomes were classified into pathovars based on the detection of specific pathovar-defining virulence factor genes by blastn. The heatmap shows the copy number of *pasA*, *pasB*, *pasC*, *pasD1*, and *pasD2* sRNA genes in the genome of each isolate detected by blastn. The six most highly represented *E. coli* clonal complexes in the ‘pas_1362’ dataset are highlighted.

Pas sRNA distribution patterns appear to correlate more closely with the pathovar identity (colored dots) of a given isolate than its phylogenetic relationship as defined by MLST-based clonal complex assignment (**Fig. 3**). For example, within the ST10 complex, all isolates classified as ‘Other’ or EAEC only encode PasB, while the EPEC isolates in this complex lack PasB and encode a variable set of PasA, PasC and PasD. This pattern of “Other” and EAEC isolates encoding only PasB and EPEC and EHEC strains encoding variable sets of PasA, PasC, and PasD is observed for all branches of the tree.

The phylogenetic relationships of ‘pas_1362’ isolates also reveal differing evolutionary paths for Pas sRNA sequence variants. While PasD1 and PasD2 can be found alone (ST11 and ST32) or together (ST29) within a given clade, and the pattern of PasD variants present is generally uniform throughout a clonal complex (**Fig. 3**). This pattern suggests that the different PasD sequence variants were acquired independently by different lineages and did not evolve *de novo* on the bacterial chromosome. Similarly, multiple independent lineages encode multiple PasA variants (**Supplementary Fig. S4**). Interestingly, PasA variants are often encoded in multiple copies, but different clades have a higher number of one specific variant; PasA is more prevalent in ST29 isolates while PasA-2 is more prevalent in ST11 isolates. Rarer sequence variants such as PasA-3, in contrast, are found only within specific clades (ST32 clonal complex) accompanied by more sequence variants (**Supplementary Fig. S4**), which may indicate that these rarer variants evolved *de novo* within these lineages.

Collectively, the distribution pattern of Pas sRNA genes within clades tracks more closely with pathovar identity rather than the phylogenetic history of an isolate. These data support a model in which repertoires of Pas sRNA genes are horizontally acquired and maintained in certain lineages of pathogenic *E. coli* rather than vertically transmitted. Notably, several well-studied non-pathogenic *E. coli* strains such as the laboratory strain K-12 and the human commensal strain HS do not contain any Pas genes (**Supplementary Table S3**). Gains or losses of individual Pas genes and sequence variants can be observed within highly clonal clades, suggesting that these genes remain subject to genetic events such as loss, mutation, duplication, or the movement of mobile genetic elements.

### Syntenic context suggests a common bacteriophage origin for certain Pas sRNAs

Local gene synteny (i.e. the order and orientation of genes in a given locus) can provide evidence of evolutionary events like horizontal gene transfer or rearrangements. To better understand the origin of Pas sRNA genes in bacterial lineages, we examined the local synteny of each Pas sRNA gene in the ‘pas_1362’ dataset. We extracted 10 kbp of flanking DNA sequence on either side of each Pas sRNA gene to generate a library of ∼20 kbp sequences that were then clustered on the basis of DNA sequence similarity using MMseqs2 [46]. The DNA sequences were then annotated with NCBI gene annotations and visualized using the R package geneviewer v.0.1.11 for manual comparison.

All five Pas sRNA genes are encoded within phage-like operons (**Fig. 4A**), consistent with their discovery in lambdoid prophage regions in *E. coli* strain E2348/69 [24]. Between 95.8-99.8% of Pas sRNA hits were found adjacent to genes encoding phage capsid or tail structural proteins (**Fig. 4B**). Additionally, transposable elements and tRNA genes, both of which are enriched in bacteriophage genomes, were also found proximal to certain Pas sRNA genes.

**Figure 4.**
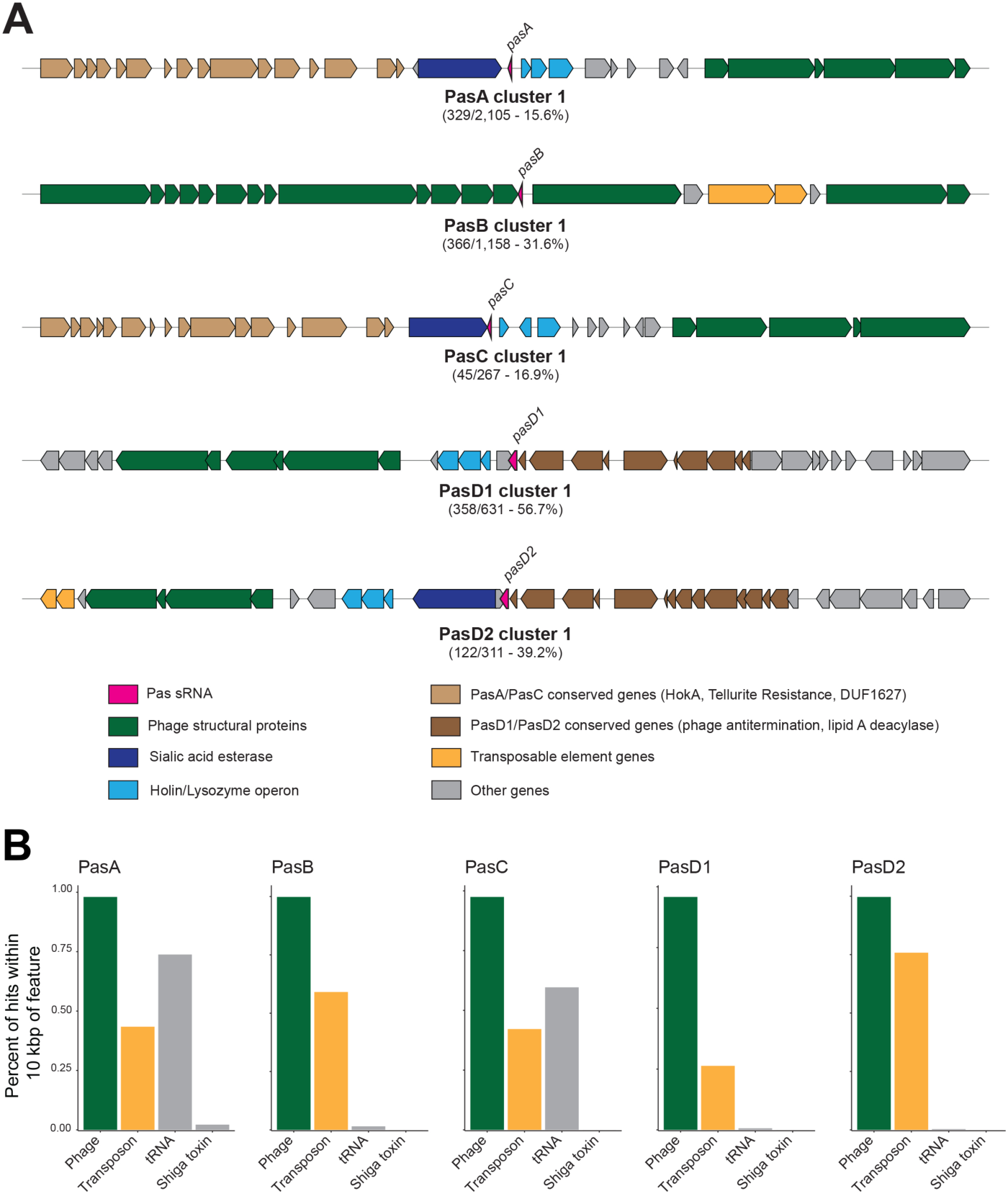
Syntenic analysis of Pas gene flanking regions by sequence clustering. (**A**) DNA sequences comprising 10 kbp flanks on either side of each *pas* gene hit were clustered using *mmseqs2* and numbered in order of prevalence. A synteny diagram for one representative region for the largest cluster (i.e. “cluster 1”) for each Pas gene is shown. Feature names and boundaries were taken from NCBI annotation data associated with each genome. (**B**) Bar graphs showing the prevalence of Pas sRNA hits within 10 kbp of genes encoding phage structural proteins (‘Phage’), transposons and transposases (‘Transposon’), tRNAs, and shiga toxins.

The regions flanking PasA and PasC showed the greatest degree of diversity, as the ∼20 kbp fragments grouped into a greater number of smaller-sized clusters compared to PasB and PasD fragments (**Supplementary Fig. S5**). The largest clusters for PasA and PasC included less than 20% of the total fragments for each gene, while the largest cluster for PasD1 included 56.7% of all PasD1 fragments. The sequence diversity in the regions surrounding PasA and PasC suggests that those loci have been subject to a greater number of evolutionary events such as recombination or horizontal transfer.

The most common gene arrangements surrounding PasA and PasC (i.e. “Cluster 1”) share a set of ‘PasA/PasC conserved genes’: RusA endonuclease (PF05866), Tellurite resistance protein TerB (PF05099), and a DUF1627-containing protein (PF07789) (**Fig. 4A**). We tabulated the frequency of these genes based on Genbank annotation files, and found this set of ‘PasA/PasC conserved genes’ was within 10 kbp of 76.8% of PasA hits and 67.6% of PasC hits (**Supplementary Table S4**). Similarly, based on the similar gene arrangements for PasD1 (Cluster 1) and PasD2 (Cluster 1), we defined a set of ‘PasD1/PasD2 conserved genes’: phage antitermination protein Q (PF03589) and lipid A deacylase LpxR (PF09982). The ‘PasD1/PasD2 conserved genes’ were found within 10 kbp of >98% of PasD1 and PasD2 hits (**Supplementary Table S4**). The overlap in neighboring genes near the *pasA*/*pasC* genes (tan arrows) as well as near the *pasD1*/*pasD2* genes (brown arrows) provide further evidence that the two pairs are sequence variants of the same sRNA gene.

The ‘PasA/PasC conserved genes’ were essentially absent (<0.05% prevalence) from the regions surrounding PasB and PasD1. However, PasA/PasC conserved genes’ are found within 10 kb of 53.9% of PasD2 hits, and ‘PasD1/PasD2 conserved genes’ are found within 10 kbp of 49.8% of PasA hits and 33.1% of PasC hits, suggesting that within certain genomes the PasD2 variant has become genetically linked to the PasA/PasC locus (**Supplementary Table S4** and **Supplementary Fig. S6A**). On the E2348/69 chromosome, PasA and PasD2 are located in the same prophage region on either side of a sialic acid esterase gene (PF03629). This arrangement of PasA and PasD2 is found 284 times in the ‘pas_1362’ dataset (**Supplementary Fig. S6B**). In the most common PasD2 gene arrangement, the sialic acid esterase gene is encoded between *pasD2* and a conserved Holin/Lysozyme operon that is typically located directly adjacent to *pasA*, *pasC*, and *pasD1* (**Fig. 4A**). Homologous recombination between similar prophage regions such as around the sialic acid esterase gene may serve as a mechanism to create genetic linkage between MGE-encoded sRNA genes. The activity of adjacent transposase genes may have contributed to the genetic linkage between PasA and PasD. The majority (75.8%) of *pasD2* genes are within 10 kbp of a transposase gene and may have become linked to *pasA* through transposition (**Supplementary Fig. S6C**).

In a parallel approach using PHASTER [47], we identified 3,343 distinct Pas gene-containing prophage regions in 1,250 bacterial genomes from the ‘pas_1362’ dataset [47]. Genomes with incomplete GenBank annotations or other errors were excluded from the analysis. With some exceptions, particularly for PasB, nearly all Pas sRNA genes mapped to intact prophage regions on bacterial chromosomes (**Supplementary Fig. S6D**). Even Pas genes located outside of PHASTER-annotated prophages are surrounded by phage genes but may lack other features recognized by PHASTER in the surrounding region. Collectively, the genetic context of Pas sRNA genes in *E. coli* genomes suggests that the genes were acquired by integration of related, lysogenic bacteriophages and have diversified through a combination of recombination and transposition events.

### Variation and the presence of multiple copies is driven by repeated acquisition

To further examine the mechanisms driving synteny variation in Pas sRNA gene loci, we examined the regions surrounding PasA genes, which show the greatest degree of both sequence and synteny diversity (**Supplementary Fig. S5**). We also investigated why PasA genes are found in higher copy numbers within genomes compared to other Pas sRNA genes. Multiple copies of *pasA* are found in 85.0% of the pasA-containing genomes. Genomes that encode multiple *copies* of *pasA* typically (87.4%) encode just PasA and PasA-2, supporting the hypothesis that the variants possess non-redundant regulatory functions. The average distance between *pasA* gene copies is nearly 0.5 Mbp, and the closest pair is still 22 kbp apart, indicating that *pasA* copies are not a product of tandem duplication (**Supplementary Fig. S7A**). In genomes that contain both *pasD1* and *pasD2* (examined collectively as ‘*pasD*’), multiple copies of *pasD* are separated by over 1.5 Mbp on average (**Supplementary Fig. S7B**).

Since there is a peak of *pasA* copies within ∼0.1 Mbp of each other, we asked whether *pasA* genes were encoded within the same prophage regions. We used PHASTER to identify the boundaries of prophage regions for chromosomes in the ‘pas_1362’ dataset and mapped the Pas sRNA genes within each prophage [47]. This analysis showed that, although some prophage regions (163) have two copies of PasA (**Supplementary Fig. S7C**), most prophage regions (2,751) contain a single Pas sRNA gene. Thus, the high copy number of *pasA* genes in bacterial genomes is driven predominantly by repeated acquisition of multiple *pasA*-containing prophage regions on the same chromosome (**Supplementary Table S3**). The reason that genomes maintain multiple copies of PasA remains unclear.

### Flanking regions encode other structured RNAs

We also searched for known conservative structural elements (CSEs) and noncoding RNAs hits located in the 10-kbp flanking regions using Infernal software [33] (cmsearch and cmscan programs) and the Rfam database [34] (see Materials and Methods for details). Based on this analysis, the most frequent CSE/ncRNA hits in the regions flanking PasA and PasC were tRNA hits (**Fig. 5A**), consistent with the MMseqs [46] analysis and the known role of tRNAs as integration sites or hot spots for prophage insertions [50]. Notably, the distribution of tRNA hits is highly similar on the minus strands of PasA and PasA-2 and closely resemble the distribution for PasC (**Supplementary Table S5**).

**Figure 5.**
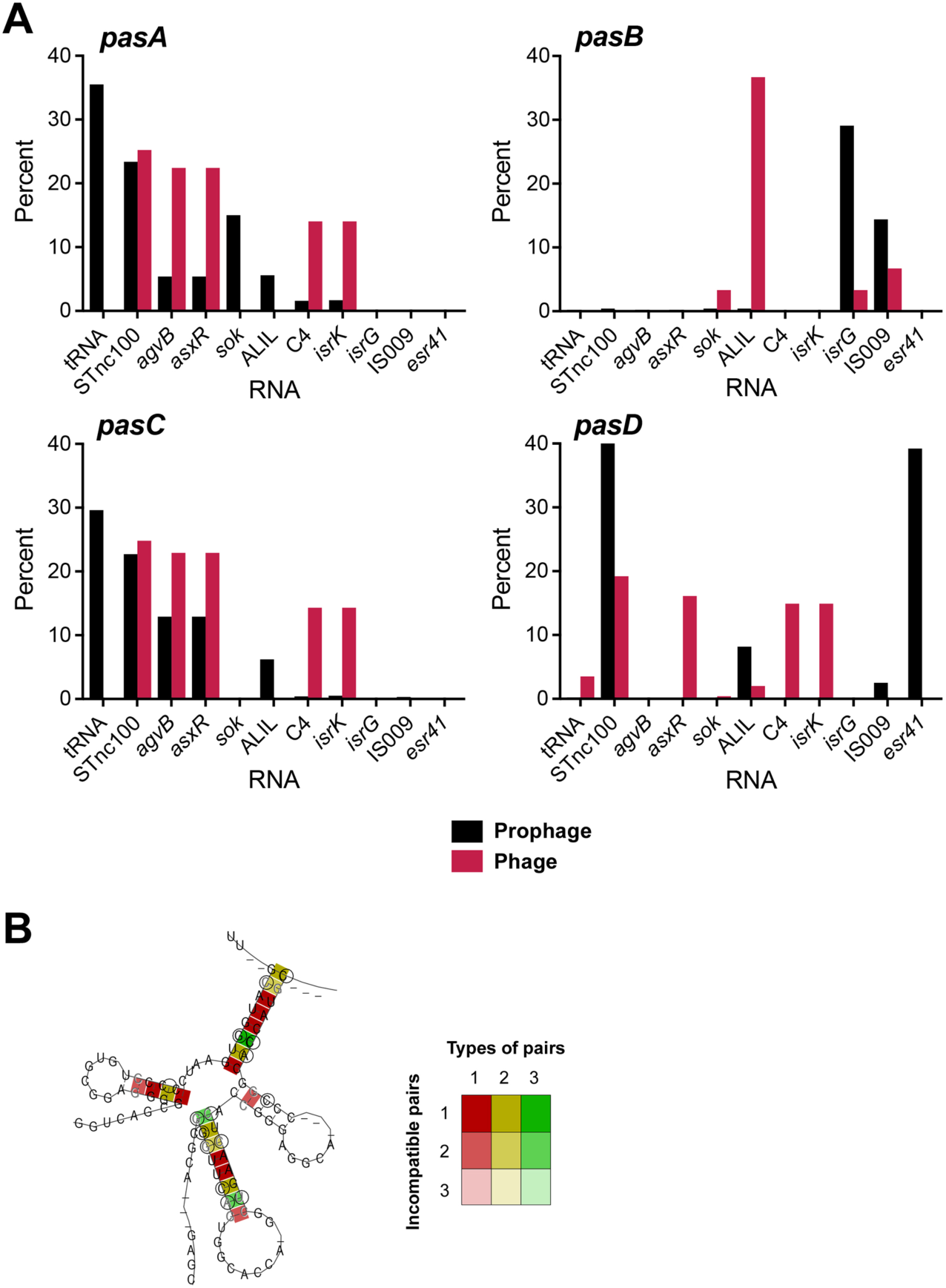
Prevalence of other sRNAs in proximity to the Pas genes in prophage and bacteriophage genomes. (**A**) Bar graphs showing the prevalence of CSE sRNA hits within 10 kbp of Pas genes in prophage and bacteriophage genomes. Only sRNAs for which the CSE hits were present as >5% of the total for at least one example are shown. Full data set is in **Supplementary Table S5**. (**B**) Predicted consensus secondary structure for MSAs of PasD and STnc100 together. RNAs were analyzed using RNAalifold [29]. Colors represent sequence variation based on the MSAs of both sRNAs (**Supplementary Fig. S8**) as described in Figure 1.

Another abundant structural hit in the PasA, PasC, and PasD flanks was STnc100 (**Fig. 5A**). The presence of this hit in the flanking regions of most PasA, PasC, and PasD sRNAs, different from the distribution of structural hits in the vicinity of PasB, is consistent with previous assertions about the diversification of gene acquisitions. The Sok RNA was also prevalent for PasA, while AgvB and AsxR were prevalent for PasC, and Esr41 was frequently found in proximity to PasD (**Fig. 5A**). Again, the PasB flanking regions displayed a distinct pattern with the most prevalent CSEs being IsrG and IS009. The STnc100 (RF02076) [51,52], Sok (RF01794) [53], AgvB (RF02702) [54], AsxR (RF02703) [54], Esr41 (RF02676) [23,55], IsrG (RF01390) [56], and IS009 (RF02111) [57] sRNAs all have previously been associated with pathogenicity islands and/or bacteriophage sequences.

During the structural searches using cmscan, we identified structural homology between PasD and STnc100 despite the sequences not being sufficiently similar such that STnc100 could be identified by BLAST searches with the PasD sequence from *E. coli* E2348/69 (**Fig. 5B**). Although the PasD and STnc100 sequences have different seed regions described to date and can interact with distinct sets of targets, it is likely that these genes originated from a common ancestor. The structural similarity between PasD and STnc100 together with associations of both PasA and PasD with esterase genes, may partially explain the enrichment of STnc100 hits in the vicinity of PasA and PasC.

### Pas sRNA sequences in bacterial genomes have diverged from bacteriophage genes

Pas sRNA genes in bacterial genomes likely originate from integrated bacteriophages, so we sought to compare the sRNA genes in prophage regions against those in sequenced bacteriophage genomes. We performed a blastn search of the five Pas sRNA genes (PasA, PasB, PasC, PasD1, and PasD2) against a set of 36,687 bacteriophage genomes available from the INPHARED data set (https://millardlab.org/phage-genomes-may-2025/) [58] (**Fig. 6A**). We also searched the same dataset for structurally similar RNAs using cmscan [33]. After filtering (e-value < 0.005 by both methods; hits > 75% query length), we identified 50 hits across 43 unique bacteriophage genomes (**Supplementary Table S6**). PasA was the most prevalent Pas sRNA found in bacteriophage genomes (25), followed by PasB (24). Two bacteriophages encoded a PasD variant that was not 100% identical to either PasD1 or PasD2 (thus we will refer to the variant as just ‘PasD’) (**Fig. 6B**). Only a single hit with 100% nucleotide identity to PasC was identified, but this because most PasC “variants” were classified as PasA due to the sequence similarity between PasA and PasC. Of the 43 Pas sRNA-encoding bacteriophages, 32 also encode Shiga toxin (Stx-converting bacteriophages) (**Fig. 6B, Supplementary Table S6**),

**Figure 6.**
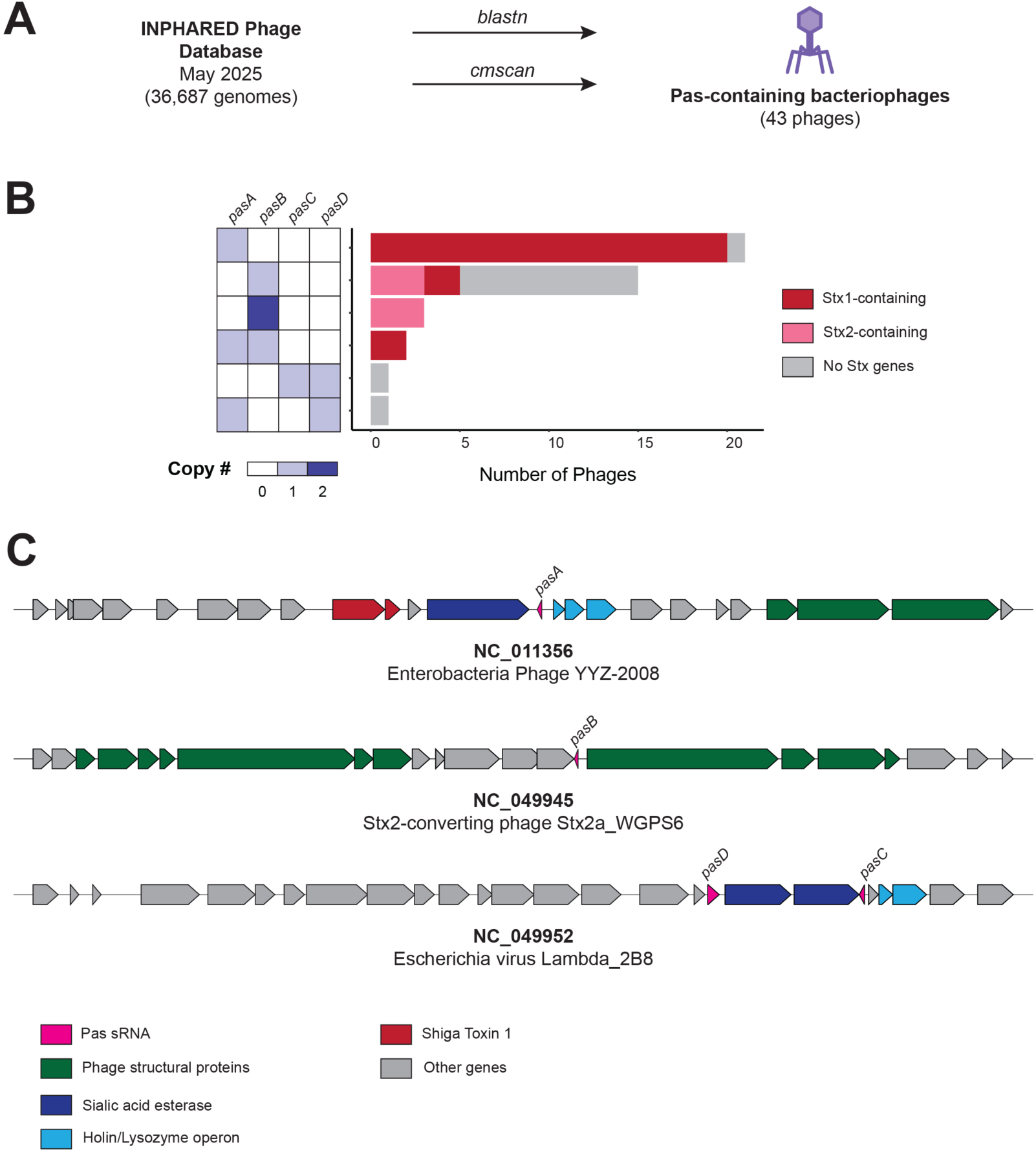
Comparison of Pas-containing bacteriophage genomes. (**A**) Simplified summary of computational approach to identifying and comparing Pas sRNA gene containing bacteriophage and prophage sequences. Bacteriophage genome sequences were identified by a blastn search of five Pas sRNA gene sequences against the INPHARED database (March 2025 version) [59,60]. BLAST searches were performed with default settings, and results were filtered for hits with >75% query coverage. (**B**) Distribution of Pas sRNA gene repertoires across 43 bacteriophage genomes containing at least one Pas sRNA. Bars are colored based on the presence of *stx1* or *stx2* Shiga toxin genes in the bacteriophage genomes. (**C**) Synteny diagrams showing 10 kbp flanking regions on either side of *pas* gene hits in the three Pas sRNA-containing NCBI RefSeq bacteriophage genomes identified by BLAST. Gene boundaries are displayed based on NCBI RefSeq annotations. In the NC_049952 annotation, the sialic acid esterase gene is split into two smaller annotated genes.

Three of the identified bacteriophages (NC_011356, NC_049945, NC_049952) are within the NCBI RefSeq database and collectively contain hits for PasA, PasB, PasC, and PasD, so they were selected as representative sequences to examine the genetic locus surrounding each Pas sRNA gene (**Fig. 6C**). In the bacteriophage genomes, the region surrounding PasB largely resembles the locus on the bacterial chromosome, where PasB is embedded within an operon of phage structural proteins (**Fig. 4A**). In contrast, the synteny of the genetic loci surrounding *pasA*, *pasC*, and *pasD* in the bacteriophage genomes differs from the bacterial chromosoms. Although the sialic acid esterase, holin, and lysozyme genes immediately flanking *pasA* and *pasC* are the same as the flanking genes on the bacterial chromosome, the region beyond the genes immediately flanking PasA and PasC is not conserved (**Fig. 6C**, **Fig. 4A**). In the representative PasA-containing phage NC_011356, a Shiga toxin 1 operon is located directly upstream of the sialic esterase gene flanking *pasA*, and 92% (22/24) of PasA-containing bacteriophage genomes also encode *stx1A*. This genetic linkage is largely absent in bacterial genomes as <3% of PasA genes on bacterial chromosomes are adjacent to Shiga toxin genes (**Fig. 4**), suggesting that genetic rearrangements have repositioned the sialic acid esterase-PasA-holin-lysozyme operon in bacterial genomes or that the *stx* gene has been repeatedly lost in bacterial genomes following horizontal acquisition. In the representative bacteriophage genome NC_049952, *pasD* is located directly upstream of the sialic acid esterase gene, which is flanked by a *pasC* gene and the holin-lysine operon. While the *pasD*-sialic acid esterase-holin-lysozyme arrangement is seen in many *E. coli* genomes (**Supplementary Fig. S6**), *pasD* variants are predominantly found alone (**Fig. 4A**). These data would suggest that PasA or PasC was lost from these PasD-containing prophage regions following their integration into bacterial chromosomes.

A comparative analysis of the 10-kbp flanks of Pas sRNAs in prophage and bacteriophage sequences revealed some significant differences. In particular, unambiguous tRNA hits were abundant only in prophage regions and could not be detected in bacteriophage genomes (**Fig. 5A**).These results are consistent with tRNA hits being considered insertion hot sports in host genomes [50]. In contrast, STnc100 hits were found very frequently in the vicinity of PasA, PasC and PasD sRNAs in both prophage and bacteriophage regions (**Fig. 5A**). Given that the most common neighboring genes for STnc100 genes in prophage regions are the putative Fels-1 prophage chaperone and Fels-1 prophage hypothetical protein, STnc100 hits in the sRNA flanking regions also can be considered as markers of the bacteriophage origin of these sequences.

Our analysis supports a model in which pathogenic *E. coli* genomes acquired PasA, PasC, and PasD sRNA genes through the integration of Stx-converting bacteriophages, a major driver of horizontal gene transfer in enterohemorrhagic and enteropathogenic lineages. Stx phages are well known to integrate at specific chromosomal attachment sites, including those within or adjacent to tRNA genes—via site-specific recombination mediated by phage-encoded integrases (reviewed in [61]). These integration hotspots promote repeated insertion and excision events and frequently result in mosaic prophage architectures. The association of Pas sRNAs with tRNA-like sequences in our datasets is consistent with this integration mechanism and suggests that these loci represent remnants of prophage insertion sites that have undergone subsequent rearrangements. PasB sRNA genes appear to originate from distinct set of bacteriophages, only some of which are Stx-converting bacteriophages, explaining their weaker association with other Pas sRNA genes and enteric pathovars of *E. coli* (**Fig. 3**). Furthermore, the syntenic conservation of the PasB flanking region on bacteriophages and in bacterial genomes contrasts with the mosaicism surrounding PasA, PasC, and PasD loci (**Supplementary Fig. S5**). Collectively, our study suggests a diversity of evolutionary paths for horizontally-acquired sRNAs in pathogenic strains of *E. coli*.

## Discussion

Here, we characterize the distribution and evolution of five sRNAs named PasA, PasB, PasC, PasD1, and PasD2 that were recently discovered in the enteropathic *E. coli* O127:H6 strain E2348/69 [24]. We show that the Pas sRNAs are only found within the *Escherichia* and *Shigella* genuses but display variability both in presence and sequence across *E. coli* strains (**Fig. 1**). Our study, in agreement with the studies in *Pseudomonas* and *Vibrio* [11,12], reinforces the idea that bacterial lineages encode sets of core sRNAs conserved at the genus-level (like OxyS) or beyond (like RyhB) along with variable sRNAs that diverge at the species- and strain-level (like the Pas sRNAs).

### Strain-level sRNA diversity can be used to inform evolutionary hypotheses

Our results highlight the utility of sampling RNAs among bacterial strains within the same species or genus for comparative studies of non-coding RNAs. The Pas sRNAs are lineage-specific and would have been missed from 88% of *E. coli* genomes available on NCBI including prominent laboratory strains such as *E. coli* K-12. Clearly, the sets of variable sRNAs within an *E. coli* genome differs from strain to strain. A previous evolutionary study attributed the absence of several *E. coli* K-12 variable sRNA genes from the E2348/69 genome to repeated gene loss and host-specific niche adaptation [44]. However, Pas sRNAs are broadly distributed across multiple *E. coli* lineages including phylogroup A, which includes *E. coli* K-12 (reviewed in [20]), and were more likely gained independently (**Fig. 3**). Thus, the sRNA gene set within any single strain may not be the ancestral state from which to infer gene gain or loss.

The diversity in Pas sRNA sequences across strains exemplifies several proposed mechanisms of sRNA evolution, including single-nucleotide polymorphisms (SNPs), genomic rearrangements, and gene duplications (reviewed in [17,62]). Most of the differences between Pas variants are SNPs though some PasA variants display larger sequence differences that likely affect the processing or existence of the sRNA. For example, PasA-3 is identical to PasA except that it has lost its Rho-indedendent terminator. Genetic rearrangements have been shown to create or destroy sRNA genes in enteric bacteria by repositioning transcriptional features [18].

The information from strain-level sRNA variance can be leveraged to infer functional information. For example, SNPs are prevalent across Pas sRNA variants but are generally not found within putative seed sequences and key structural features like Rho-independent terminator loops (**Fig. 1**). Previously unknown features, such as a putative seed sequence in PasB (**Supplementary Fig. S1**), could be suggested by identifying regions of sequence conservation. For each Pas sRNA, several sequence variants that differ by only a few SNPs have become fixed within the population, suggesting that these variants may be undergoing selection. Although these SNPs typically only involve a few sites, they may result in structural difference across Pas variants (**Supplementary Fig. S2**). Minor sequence changes in sRNAs have been shown to alter protein binding and regulatory function [63]. Indeed, PasD1 and PasD2 differ by 4 SNPs but bind a different set of targets in E2348/69 [24].

Sequence variants of PasA and PasD are often encoded in duplicate within the same genome. However, unlike other examples of duplicated sRNAs such as OmrA/B or PrrF1/2 in *Pseudomonas* [64,65], multiple copies of PasA and PasD are encoded distally and were likely acquired by superinfection with multiple Pas sRNA-carrying bacteriophages rather than by tandem duplication. The two most prevalent PasA variants, PasA and PasA-2, are frequently found together within a genome, but the copy number (0-5) of each variant differs from one strain to another (**Supplementary Fig. S4**), reminiscent of the quorum sensing Qrr sRNAs in *Vibrio* [66]. PasA and PasA-2 share a conserved seed sequence, but the 5’ sequence is largely divergent (**Fig. 1**), which may lead to differences in processing or expression between PasA and PasA-2. The RNA-RNA interactome of PasA-2 is unknown, because this sRNA is absent from the E2348/69 genome in which the RIL-seq analysis for PasA was performed [26]. PasD1 and PasD2, the two most prevalent sequence variants of PasD, are both found in E2348/69 and share an overlapping but distinct interactome [24]. The variants are found individually in certain *E. coli* lineages and together in others. Loss of one PasD variant appears to be more common in lineages that encode both PasD1 and PasD2, such as ST29, which may indicate some redundancy in their function (**Fig. S4**). Further experimental investigation of functional differences between PasA and PasD sequence variants is required to discern whether the duplication of sequence variants represents the bona fide evolution of separate sRNAs or reflects a relatively young sRNA gene still undergoing selection. Gene duplication through repeated horizontal transfer uniquely affects accessory genome sRNAs and could serve as an evolutionary mechanism of generating neofunctional regulators.

In our sequence comparisons, we noted that PasA variants can be aligned with PasC variants (**Supplementary Fig. S1B**) indicating a common origin for these two sRNAs. We could similarly align PasD variants with the STnc100 sRNA, again suggesting that these two families had a common origin. We note that these similarities mean that for both of the pairs, some variants mistakenly could be annotated as a member of the other family. Finally, we detected genetic linkage between the PasD2 variant and PasA/PasC family members, though the physiological advantage of this linkage is not known.

### Pas sRNAs originate from Stx-converting phages but have greatly diverged

Large portions of the genome in pathogenic *E. coli* strains originate from phages [67], and genomes in the ‘pas_1362’ dataset have an average of 17 prophage regions. Based on their genetic context, Pas sRNAs were almost certainly acquired in bacterial genomes through the integration of lysogenic phages. A BLAST search for Pas genes in the NCBI ‘Complete_Bacteriophage’ database revealed seven bacteriophage genomes that encoded homologs of Pas sRNAs. However, a majority of the Pas-containing prophage regions on bacterial chromosomes in the ‘pas_1362’ dataset resemble other phages that do not encode Pas genes in their sequenced genomes. Furthermore, with the exception of PasA, the nucleotide sequence of Pas homologs in the bacteriophage genomes are divergent from those in bacteria.

The paucity of Pas-containing bacteriophage genomes could reflect a limitation in the number of sequenced bacteriophage genomes available. However, an additional BLAST search for Pas sRNA genes in a database of 189,680 viral genomes assembled from human stool metagenomes in 2021 yielded no hits [27]. This would suggest that the population of extant bacteriophages have evolutionarily diverged from the pool of bacteriophages that originally introduced Pas sRNAs into bacterial genomes. Alternatively, the matching of prophage regions with bacteriophage genomes may be obscured by genetic rearrangements between homologous phage sequences. All of the phages identified by BLAST or PHASTER are lambdoid phages in the class Caudoviricetes, and several are related Shiga toxin-encoding bacteriophages (Stx phages). Stx phage genomes are mosaic due to recombination events with host genes and coninfecting phages [68,69]. The mean protein similarity between prophage regions in the ‘pas_1362’ dataset and any single bacteriophage was only 32.3%. Thus, the prophage regions are likely chimeras of multiple phages.

Estimating the age of horizontally transferred sRNAs is difficult, because different lineages of bacteria may have acquired the same MGE at different times. PasA and PasB are found most distantly in *Escherichia albertii* and related *Shigella boydii* strains, which diverged from *E. coli* roughly 28 million years ago [70]. However, the sequences of Pas genes in *E. albertii* and *Shigella* are identical to those in *E. coli* and were probably acquired by phage after the phylogenetic split. Most likely, Pas sRNA genes emerged in parallel with the Stx phages within the last 5 million years [71]. Modern *E. coli* lineages are still undergoing pathogenic conversion by Stx phages and may have acquired Pas sRNAs very recently. For example, *E. coli* strains in the pathogenic ST29 clonal complex, which emerged in the 20^th^ century [72], encode multiple copies of *pasA* and *pasD* variants that are variably gained or lost across strains (**Supplementary Fig. S4**), suggesting that they are still undergoing selection. Thus, the distribution of Pas sRNAs may reflect the age of Stx phages in a lineage.

Previous studies comparing core and variable sRNAs in *E. coli* identified several metrics correlated with sRNA age: younger sRNAs are expressed at lower levels, are less well integrated into regulatory networks, and show higher levels of sequence diversity [7,10]. In comparison to other novel sRNA candiates, the Pas sRNAs are all expressed at high levels and involved in numerous RNA-RNA interactions in E2348/69, so they appear to be relatively old within this strain [24]. However, the characteristics of Pas sRNA expression may differ in other strains. Among the Pas sRNAs, PasA and PasD show the greatest degree of sequence and synteny diversity (**Supplementary Fig. S5**), while, PasB shows the lowest degree of sequence and synteny diversity, suggesting that it is the oldest of the Pas sRNAs. Concordantly, PasB is more frequently located “outside” of prophage regions on the chromosome (**Supplementary Fig. S6D**), which may be evidence of neighboring phage genes degrading over evolutionary time.

### Possible plasticity in role of sRNAs encoded by core genome, prophage or phage

The physiological impact of sRNA regulation within a cell depends on the repertoire of sRNAs, target genes, and RNA-binding proteins present within a cell, and the effects of manipulating a single element may differ depending on the context. Although there is a clear association between Pas sRNA genes and pathogenicity in *E. coli*, the presence of Pas sRNAs is not a universal feature of pathogenic strains (**Fig. 2**), possibly due to differences in the target mRNA repertoire. It is thought that novel sRNAs are lost from genomes unless they can integrate into existing regulatory networks [7], and new sRNAs precede the emergence of target binding sites. Comparative analysis has shown that some target binding sites in enteric bacteria are not conserved even in genomes that encode both the sRNA and its target gene and for which regulation has been documented in a different strain [9,73]. Like many other horizontally acquired sRNAs (reviewed in [19]), the Pas sRNAs in E2348/69 interact with numerous genes in both the core *E. coli* genome and virulence factor genes encoded on other horizontally acquired elements. To gain further insights into the co-evolution of Pas sRNAs and their regulatory targets, experimental validation of target binding sites will need to be carried out in more strains.

Our results demonstrates the power of using publicly available bacterial sequence data for comparative studies of RNA evolution. Investigating the function of sRNAs in bacterial pathogens remains important, as post-transcriptional regulation by bacterial sRNAs has been increasingly implicated in virulence and antimicrobial resistance (reviewed in [74,75]). Ultimately, the Pas sRNAs from E2348/69 represent only a fraction of the sRNA content in the *E. coli* pangenome. Numerous other variable sRNAs have been documented in the accessory genome of other pathogenic *E. coli* strains such as EHEC O157:H7 strain Sakai [54]. Different bacterial strains from different niches likely harbor their own set of undiscovered sRNAs. The information gained from comparative studies can help guide *in silico* approaches for sRNA prediction [76] and create a framework for future experiments to dissect the relationship between sRNA-based regulation and bacterial pathogenesis.

## Acknowledgements

We thank John Dekker and Grace Morales for critical reading and comments. This work utilized the computational resources of the NIH HPC Biowulf cluster (https://hpc.nih.gov).

This research was supported by the Intramural Research Program of the National Institutes of Health (NIH). The contributions of the NIH authors are considered Works of the United States Government. The findings and conclusions presented in this paper are those of the authors and do not necessarily reflect the views of the NIH or the U.S. Department of Health and Human Services.

## Author contributions

Dennis Zhu (Conceptualization [lead], Data curation [lead], Formal analysis [equal], Investigation [equal], Methodology [equal], Visualization [lead], Writing—original draft [lead], Writing—review & editing [equal]), Svetlana A Shabalina (Data curation [supporting], Formal analysis [equal], Investigation [equal], Methodology [supporting], Visualization [supporting], Writing—original draft [supporting], Writing—review & editing [supporting]), and Gisela Storz (Conceptualization [supporting], Visualization [supporting], Writing—original draft [supporting], Writing—review & editing [equal]).

## Supplementary data

Supplementary data is available at NAR online.

## Data availability

The data underlying this article are available in the article and in its online supplementary material.

## References

1. Wagner, E.G.H. and Romby, P. (2015) Small RNAs in bacteria and archaea: who they are, what they do, and how they do it. Adv. Genet., 90, 133–208.

2. Storz, G., Vogel, J. and Wassarman, K.M. (2011) Regulation by small RNAs in bacteria: expanding frontiers. Mol. Cell, 43, 880–891.

3. Papenfort, K. and Melamed, S. (2023) Small RNAs, large networks: posttranscriptional regulons in gram-negative bacteria. Annu. Rev. Microbiol., 77, 23–43.

4. (2019) Regulating with RNA in Bacteria and Archaea. ASM Press, Washington, District of Columbia.

5. Dutcher, H.A. and Raghavan, R. (2018) Origin, evolution, and loss of bacterial small RNAs. Microbiol. Spectr., 6, 10.1128/microbiolspec.rwr-0004-2017.

6. Adams, P.P. and Storz, G. (2020) Prevalence of small base-pairing RNAs derived from diverse genomic loci. Biochim. Biophys. Acta. Gene Regul. Mech., 1863, 194524.

7. Skippington, E. and Ragan, M.A. (2012) Evolutionary dynamics of small RNAs in 27 *Escherichia coli* and *Shigella* genomes. Genome Biol. Evol., 4, 330–345.

8. Lindgreen, S., Umu, S.U., Lai, A.S., Eldai, H., Liu, W., McGimpsey, S., Wheeler, N.E., Biggs, P.J., Thomson, N.R., Barquist, L. et al. (2014) Robust identification of noncoding RNA from transcriptomes requires phylogenetically-informed sampling. PLos Comput. Biol., 10, e1003907.

9. Peer, A. and Margalit, H. (2014) Evolutionary patterns of *Escherichia coli* small RNAs and their regulatory interactions. RNA, 20, 994–1003.

10. Kacharia, F.R., Millar, J.A. and Raghavan, R. (2017) Emergence of new sRNAs in enteric bacteria is associated with low expression and rapid evolution. J. Mol. Evol., 84, 204–213.

11. Toffano-Nioche, C., Nguyen, A.N., Kuchly, C., Ott, A., Gautheret, D., Bouloc, P. and Jacq, A. (2012) Transcriptomic profiling of the oyster pathogen *Vibrio splendidus* opens a window on the evolutionary dynamics of the small RNA repertoire in the *Vibrio* genus. RNA, 18, 2201–2219.

12. Trouillon, J., Han, K., Attrée, I. and Lory, S. (2022) The core and accessory Hfq interactomes across *Pseudomonas aeruginosa* lineages. Nat. Commun., 13, 1258.

13. Chareyre, S. and Mandin, P. (2018) Bacterial iron homeostasis regulation by sRNAs. Microbiol. Spectr., 6, 10.1128/microbiolspec.rwr-0010-2017.

14. Iosub, I.A., van Nues, R.W., McKellar, S.W., Nieken, K.J., Marchioretto, M., Sy, B., Tree, J.J., Viero, G. and Granneman, S. (2020) Hfq CLASH uncovers sRNA-target interaction networks linked to nutrient availability adaptation. eLife, 9, e54655.

15. Melamed, S., Peer, A., Faigenbaum-Romm, R., Gatt, Y.E., Reiss, N., Bar, A., Altuvia, Y., Argaman, L. and Margalit, H. (2016) Global mapping of small RNA-target interactions in bacteria. Mol. Cell, 63, 884–897.

16. Melamed, S. (2020) New sequencing methodologies reveal interplay between multiple RNA-binding proteins and their RNAs. Curr. Genet., 66, 713–717.

17. Updegrove, T.B., Shabalina, S.A. and Storz, G. (2015) How do base-pairing small RNAs evolve? FEMS Microbiol. Rev., 39, 379–391.

18. Raghavan, R., Kacharia, F.R., Millar, J.A., Sislak, C.D. and Ochman, H. (2015) Genome rearrangements can make and break small RNA genes. Genome Biol. Evol., 7, 557–566.

19. Fröhlich, K.S. and Papenfort, K. (2016) Interplay of regulatory RNAs and mobile genetic elements in enteric pathogens. Mol. Microbiol., 101, 701–713.

20. Denamur, E., Clermont, O., Bonacorsi, S. and Gordon, D. (2021) The population genetics of pathogenic *Escherichia coli*. Nat. Rev. Microbiol., 19, 37–54.

21. Hansen, A.-M. and Kaper, J.B. (2009) Hfq affects the expression of the LEE pathogenicity island in enterohaemorrhagic *Escherichia coli*. Mol. Microbiol., 73, 446–465.

22. Kendall, M.M., Gruber, C.C., Rasko, D.A., Hughes, D.T. and Sperandio, V. (2011) Hfq virulence regulation in enterohemorrhagic *Escherichia coli* O157:H7 strain 86-24. J. Bacteriol., 193, 6843–6851.

23. Suko, N., Soma, A., Iyoda, S., Oshima, T., Ohto, Y., Saito, K. and Sekine, Y. (2018) Small RNA Esr41 inversely regulates expression of LEE and flagellar genes in enterohaemorrhagic *Escherichia coli*. Microbiology (Reading), 164, 821–834.

24. Pearl Mizrahi, S., Elbaz, N., Argaman, L., Altuvia, Y., Katsowich, N., Socol, Y., Bar, A., Rosenshine, I. and Margalit, H. (2021) The impact of Hfq-mediated sRNA-mRNA interactome on the virulence of enteropathogenic *Escherichia coli*. Sci. Adv., 7, eabi8228.

25. Bar, A., Argaman, L., Altuvia, Y. and Margalit, H. (2021) Prediction of novel bacterial small RNAs From RIL-Seq RNA-RNA interaction data. Front. Microbiol., 12, 635070.

26. Iguchi, A., Thomson, N.R., Ogura, Y., Saunders, D., Ooka, T., Henderson, I.R., Harris, D., Asadulghani, M., Kurokawa, K., Dean, P. et al. (2009) Complete genome sequence and comparative genome analysis of enteropathogenic *Escherichia coli* O127:H6 strain E2348/69. J. Bacteriol., 191, 347–354.

27. Nayfach, S., Páez-Espino, D., Call, L., Low, S.J., Sberro, H., Ivanova, N.N., Proal, A.D., Fischbach, M.A., Bhatt, A.S., Hugenholtz, P. et al. (2021) Metagenomic compendium of 189,680 DNA viruses from the human gut microbiome. Nat. Microbiol., 6, 960–970.

28. O’Leary, N.A., Cox, E., Holmes, J.B., Anderson, W.R., Falk, R., Hem, V., Tsuchiya, M.T.N., Schuler, G.D., Zhang, X., Torcivia, J. et al. (2024) Exploring and retrieving sequence and metadata for species across the tree of life with NCBI Datasets. Sci. Data, 11, 732.

29. Lorenz, R., Bernhart, S.H., Höner Zu Siederdissen, C., Tafer, H., Flamm, C., Stadler, P.F. and Hofacker, I.L. (2011) ViennaRNA Package 2.0. Algorithms Mol. Biol., 26.

30. Edgar, R.C. (2022) Muscle5: High-accuracy alignment ensembles enable unbiased assessments of sequence homology and phylogeny. Nat. Commun., 13, 6968.

31. Ogurtsov, A.Y., Shabalina, S.A., Kondrashov, A.S. and Roytberg, M.A. (2006) Analysis of internal loops within the RNA secondary structure in almost quadratic time. Bioinformatics, 22, 1317–1324.

32. Matveeva, O.V. and Shabalina, S.A. (1993) Intermolecular mRNA-rRNA hybridization and the distribution of potential interaction regions in murine 18S rRNA. Nucleic Acids Res., 21, 1007–1011.

33. Nawrocki, E. and Eddy, S.R. (2013) Infernal 1.1: 100-fold faster RNA homology searches. Bioinformatics, 29, 2933–2935.

34. Ontiveros-Palacios, N., Cooke, E., Nawrocki, E.P., Triebel, S., Marz, M., Rivas, E., Griffiths-Jones, S., Petrov, A.I., Bateman, A. and Sweeney, B. (2025) Rfam 15: RNA families database in 2025. Nucleic Acids Res., 53, D258–D267.

35. Ogurtsov, A.Y., Roytberg, M.A., Shabalina, S.A. and Kondrashov, A.S. (2002) OWEN: aligning long collinear regions of genomes. Bioinformatics, 18, 1703–1704.

36. Kondrashov, A.S. and Shabalina, S.A. (2002) Classification of common conserved sequences in mammalian intergenic regions. Hum. Mol. Genet., 11, 669–674.

37. Tamura, K., Stecher, G. and Kumar, S. (2021) MEGA11: Molecular Evolutionary Genetics Analysis Version 11. Mol. Biol. Evol., 38, 3022–3027.

38. Kumar, S., Stecher, G., Li, M., Knyaz, C. and Tamura, K. (2018) MEGA X: Molecular Evolutionary Genetics Analysis across Computing Platforms. Mol. Biol. Evol., 35, 1547–1549.

39. Kimura, M. (1980) A simple method for estimating evolutionary rate of base substitutions through comparative studies of nucleotide sequences. J. Mol. Evol., 16, 111–120.

40. Schoch, C.L., Ciufo, S., Domrachev, M., Hotton, C.L., Kannan, S., Khovanskaya, R., Leipe, D., Mcveigh, R., O’Neill, K., Robbertse, B. et al. (2020) NCBI Taxonomy: a comprehensive update on curation, resources and tools. Database (Oxford), baaa062.

41. Tartof, S.Y., Solberg, O.D., Manges, A.R. and Riley, L.W. (2005) Analysis of a uropathogenic *Escherichia coli* clonal group by multilocus sequence typing. J. Clin. Microbiol., 43, 5860–5864.

42. Tatusova, T., DiCuccio, M., Badretdin, A., Chetvernin, V., Nawrocki, E., Zaslavsky, L., Lomsadze, A., Pruitt, K.D., Borodovsky, M. and Ostell, J. (2016) NCBI prokaryotic genome annotation pipeline. Nucleic Acids Res., 44, 6614–6624.

43. Jolley, K.A., Bray, J.E. and Maiden, M.C.J. (2018) Open-access bacterial population genomics: BIGSdb software, the PubMLST.org website and their applications. Wellcome Open Res., 3, 124.

44. Stamatakis, A. (2014) RAxML version 8: a tool for phylogenetic analysis and post-analysis of large phylogenies. Bioinformatics, 30, 1312–1313.

45. Yu, G. (2020) Using ggtree to visualize data on tree-like structures. Curr. Protoc. Bioinformatics, 69, e96.

46. Steinegger, M. and Söding, J. (2017) MMseqs2 enables sensitive protein sequence searching for the analysis of massive data sets. Nat. Biotechnol., 35, 1026–1028.

47. Arndt, D., Grant, J.R., Marcu, A., Sajed, T., Pon, A., Liang, Y. and Wishart, D.S. (2016) PHASTER: a better, faster version of the PHAST phage search tool. Nucleic Acids Res., 44, W16–21.

48. Bernhart, S.H., Hofacker, I.L., Will, S., Gruber, A.R. and Stadler, P.F. (2008) RNAalifold: improved consensus structure prediction for RNA alignments. BMC Bioinformatics, 9, 474.

49. Nicolas-Chanoine, M.-H., Bertrand, X. and Madec, J.-Y. (2014) *Escherichia coli* ST131, an intriguing clonal group. Clin. Microbiol. Rev., 27, 543–574.

50. Canchaya, C., Fournous, G. and Brüssow, H. (2004) The impact of prophages on bacterial chromosomes. Mol. Microbiol., 53, 9–18.

51. Boutet, E., Djerroud, S. and Perreault, J. (2022) Small RNAs beyond model organisms: Have we only scratched the surface? Int. J. Mol. Sci., 23, 4448.

52. Paganini, J.A., Khatun, S., McAteer, S.P., Cowley, L., Greig, D.R., Gally, D.L., Jenkins, C. and Dallman, T.J. (2025) Predicting clinical outcome of Escherichia coli O157:H7 infections using explainable machine learning. Microb. Genom., 11, 001591.

53. Takada, K., Hama, K., Sasaki, T. and Otsuka, Y. (2021) The *hokW-sokW* locus encodes a type I toxin–antitoxin system that facilitates the release of lysogenic Sp5 phage in enterohemorrhagic *Escherichia coli* O157. Toxins, 13, 796.

54. Tree, J.J., Granneman, S., McAteer, S.P., Tollervey, D. and Gally, D.L. (2014) Identification of bacteriophage-encoded anti-sRNAs in pathogenic *Escherichia coli*. Mol. Cell, 55, 199–213.

55. Sudo, N., Soma, A., Muto, A., Iyoda, S., Suh, M., Kurihara, N., Abe, H., Tobe, T., Ogura, Y., Hayashi, T. et al. (2014) A novel small regulatory RNA enhances cell motility in enterohemorrhagic *Escherichia coli*. J. Gen. Appl. Microbiol., 60, 44–50.

56. Padalon-Brauch, G., Hershberg, R., Elgrably-Weiss, M., Baruch, K., Rosenshine, I., Margalit, H. and Altuvia, S. (2008) Small RNAs encoded within genetic islands of *Salmonella typhimurium* show host-induced expression and role in virulence. Nucleic Acids Res., 36, 1913–1927.

57. Neuhaus, K., Landstorfer, R., Simon, S., Schober, S., Wright, P.R., Smith, C., Backofen, R., Wecko, R., Keim, D.A. and Scherer, S. (2017) Differentiation of ncRNAs from small mRNAs in *Escherichia coli* O157:H7 EDL933 (EHEC) by combined RNAseq and RIBOseq - *ryhB* encodes the regulatory RNA RyhB and a peptide, RyhP. BMC Genomics, 18, 216.

58. Cook, R., Brown, N., Redgwell, T., Rihtman, B., Barnes, M., Clokie, M., Stekel, D.J., Hobman, J., Jones, M.A. and Millard, A. (2021) INfrastructure for a PHAge REference Database: Identification of large-scale biases in the current collection of cultured phage genomes. Phage (New Rochelle), 2, 214–223.

59. Camacho, C., Coulouris, G., Avagyan, V., Ma, N., Papadopoulos, J., Bealer, K. and Madden, T.L. (2009) BLAST+: architecture and applications. BMC Bioinformatics, 10, 421.

60. Cook, R., Brown, N., Redgwell, T., Rihtman, B., Barnes, M., Clokie, M., Stekel, D.J., Hobman, J., Jones, M.A. and A., M. (2021) INfrastructure for a PHAge REference Database: Identification of Large-Scale Biases in the Current Collection of Cultured Phage Genomes. Phage (New Rochelle), 2, 214–223.

61. Krüger, A. and Lucchesi, P.M. (2015) Shiga toxins and stx phages: highly diverse entities. Microbiology (Reading), 161, 451–462.

62. Caswell, C.C., Oglesby-Sherrouse, A.G. and Murphy, E.R. (2014) Sibling rivalry: related bacterial small RNAs and their redundant and non-redundant roles. Front. Cell Infect. Microbiol., 4, 151.

63. Peterman, N., Lavi-Itzkovitz, A. and Levine, E. (2014) Large-scale mapping of sequence-function relations in small regulatory RNAs reveals plasticity and modularity. Nucleic Acids Res., 42, 12177–12188.

64. Guillier, M. and Gottesman, S. (2006) Remodelling of the Escherichia coli outer membrane by two small regulatory RNAs. Mol. Microbiol., 59, 231–247.

65. Wilderman, P.J., Sowa, N.A., FitzGerald, D.J., FitzGerald, P.C., Gottesman, S., Ochsner, U.A. and Vasil, M.L. (2004) Identification of tandem duplicate regulatory small RNAs in *Pseudomonas aeruginosa* involved in iron homeostasis. Proc. Natl. Acad. Sci. USA, 101, 9792–9797.

66. Lenz, D.H., Mok, K.C., Lilley, B.N., Kulkarni, R.V., Wingreen, N.S. and Bassler, B.L. (2004) The small RNA chaperone Hfq and multiple small RNAs control quorum sensing in *Vibrio harveyi* and *Vibrio cholerae*. Cell, 118, 69–82.

67. Rasko, D.A., Rosovitz, M.J., Myers, G.S., Mongodin, E.F., Fricke, W.F., Gajer, P., Crabtree, J., Sebaihia, M., Thomson, N.R., Chaudhuri, R. et al. (2008) The pangenome structure of *Escherichia coli*: comparative genomic analysis of *E. coli* commensal and pathogenic isolates. J. Bacteriol., 190, 6881–6893.

68. Brüssow, H., Canchaya, C. and Hardt, W.-D. (2004) Phages and the evolution of bacterial pathogens: from genomic rearrangements to lysogenic conversion. Microbiol. Mol. Biol. Rev., 68, 560–602.

69. Smith, D.L., Rooks, D.J., Fogg, P.C.M., Darby, A.C., Thomson, N.R., McCarthy, A.J. and Allison, H.E. (2012) Comparative genomics of Shiga toxin encoding bacteriophages. BMC Genomics, 13, 311.

70. Hyma, K.E., Lacher, D.W., Nelson, A.M., Bumbaugh, A.C., Janda, J.M., Strockbine, N.A., Young, V.B. and Whittam, T.S. (2005) Evolutionary genetics of a new pathogenic *Escherichia* species: *Escherichia albertii* and related *Shigella boydii* strains. J. Bacteriol., 187, 619–628.

71. Reid, S.D., Herbelin, C.J., Bumbaugh, A.C., Selander, R.K. and Whittam, T.S. (2000) Parallel evolution of virulence in pathogenic *Escherichia coli*. Nature, 406, 64–67.

72. Zhang, W.L., Bielaszewska, M., Liesegang, A., Tschäpe, H., Schmidt, H., Bitzan, M. and Karch, H. (2000) Molecular characteristics and epidemiological significance of Shiga toxin-producing *Escherichia coli* O26 strains. J. Clin. Microbiol., 38, 2134–2140.

73. Richter, A.S. and Backofen, R. (2012) Accessibility and conservation: general features of bacterial small RNA-mRNA interactions? RNA Biol., 9, 954–965.

74. Felden, B. and Augagneur, Y. (2021) Diversity and versatility in small RNA-mediated regulation in bacterial pathogens. Front. Microbiol., 12, 71997.

75. Mediati, D.G., Wu, S., W., W. and Tree, J.J. (2021) Networks of resistance: small RNA control of Antibiotic Resistance. Trends Genet., 37, 35–45.

76. Wright, P.R., Richter, A.S., Papenfort, K., Mann, M., Vogel, J., Hess, W.R., Backofen, R. and Georg, J. (2013) Comparative genomics boosts target prediction for bacterial small RNAs. Proc. Natl. Acad. Sci. USA, 110, E3487–E3496.

