## Supplementary Figures for "Distribution of prophage-encoded Pas sRNAs across pathogenic *Escherichia coli*"

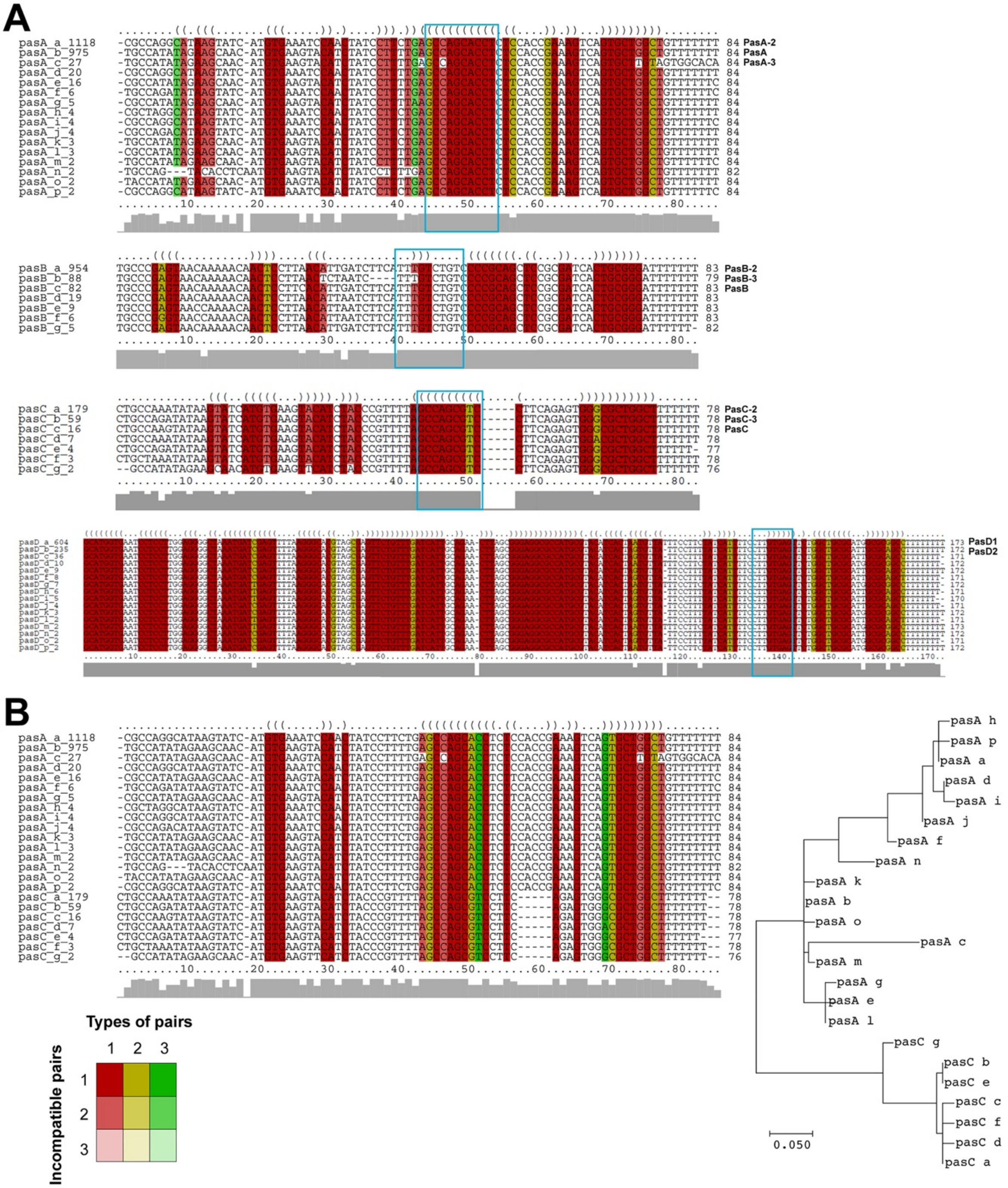


**Figure S1.** Examples of multiple alignments and profiles of sequence identity of analyzed Pas RNAs (**A**) The coloring scheme used to highlight the mutational pattern with respect to the secondary structure (folding), was is shown in the insert. If one predicted base-pair is formed by several different combinations of nucleotides, compensatory mutations have occured. This is indicated by different colors. Pale colors indicate that a base pair cannot be formed in some sequences of the alignment. Consensus folding (upper row) is indicated for a sequence of equal length consisting of matching brackets and dots. A base pair between two positions is represented by the symbol '(' at the first position and ')' at the second position, while unpaired bases are represented by dots. The identity profiles of the analyzed Pas sRNA sequences are shown at the bottom in grey. Nucleotides likely involved in intermolecular interactions (seed regions) are highlighted with blue boxes. The seed sequences reported in [1] are shown for PasA and PasD. The seed sequence for PasC was inferred from PasA based on sequence conservation and structural similarity, and the seed sequence for PasB was predicted from sequence and structural conservation. Analysis was only carried out for homologs detected in more than one strain and are labeled alphabetically with number of examples. Names for previously identified Pas RNAs and examples studied more extensively are give on the right side of the alignments. (**B**) Similarity between PasA and PasC and corresponding evolutionary tree. Coloring scheme for alignment and nomenclature is as in (A). The evolutionary history was inferred by using the Maximum Likelihood method and Kimura 2-parameter model [2]. The tree with the highest log likelihood (-393.66) is shown. Initial tree(s) for the heuristic search were obtained automatically by applying Neighbor-Join and BioNJ algorithms to a matrix of pairwise distances estimated using the Maximum Composite Likelihood (MCL) approach, and then selecting the topology with superior log likelihood value. A discrete Gamma distribution was used to model evolutionary rate differences among sites (5 categories (+G, parameter = 1.5054)). The tree is drawn to scale, with branch lengths measured in the number of substitutions per site. This analysis involved 23 nucleotide sequences. All positions with less than 50% site coverage were eliminated, i.e., fewer than 50% alignment gaps, missing data, and ambiguous bases were allowed at any position (partial deletion option). There were a total of 84 positions in the final dataset. Evolutionary analyses were conducted in MEGA11 [3].

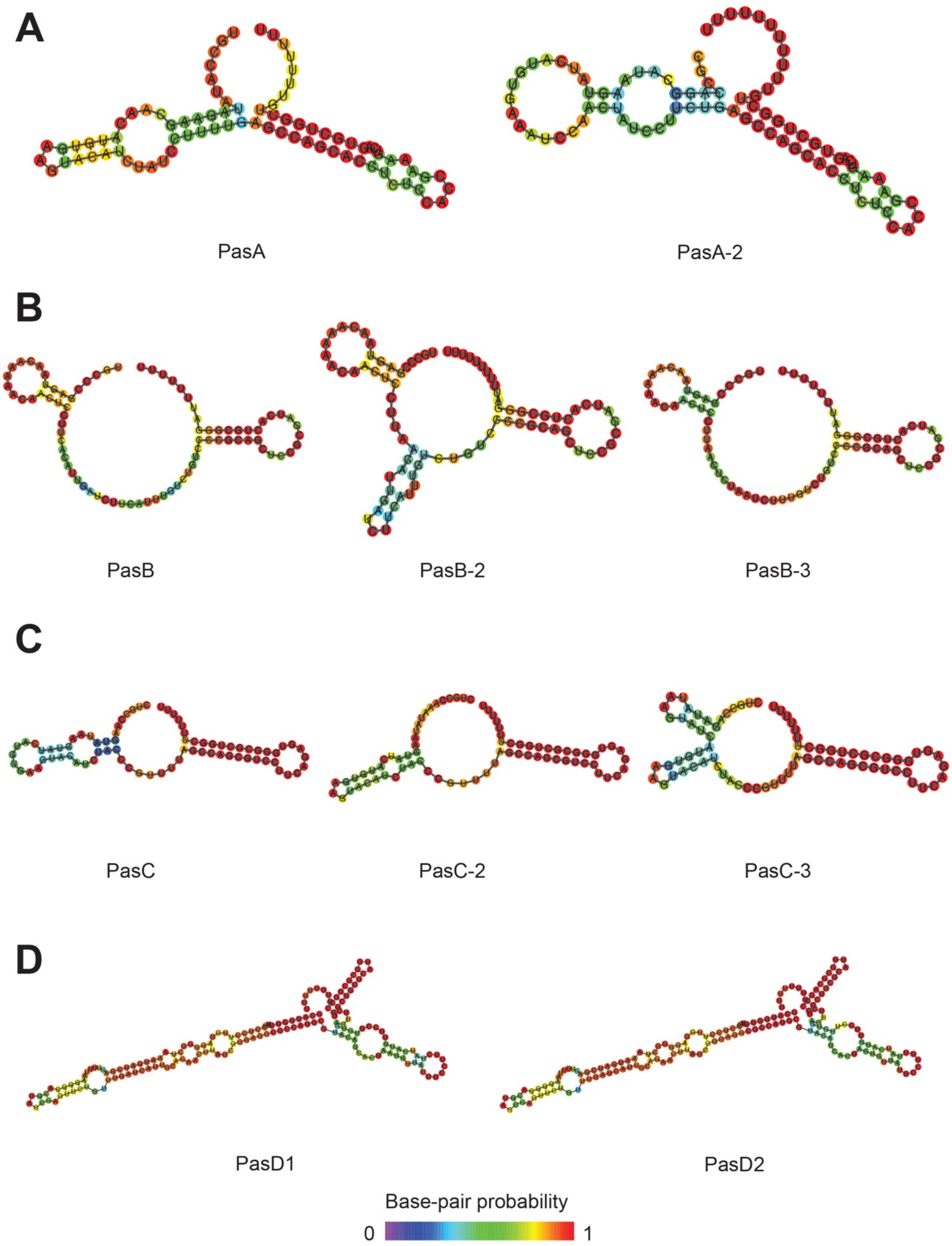


**Figure S2.** Minimum free energy RNA folding structures predicted by RNAfold software for (A) PasA variants, (B) PasB variants, (C) PasC variants, and (D) PasD1 and PasD2. Folding was performed using default parameters and folding conditions for RNAfold 2.6.3. Bases are colored by their base-pairing probability.

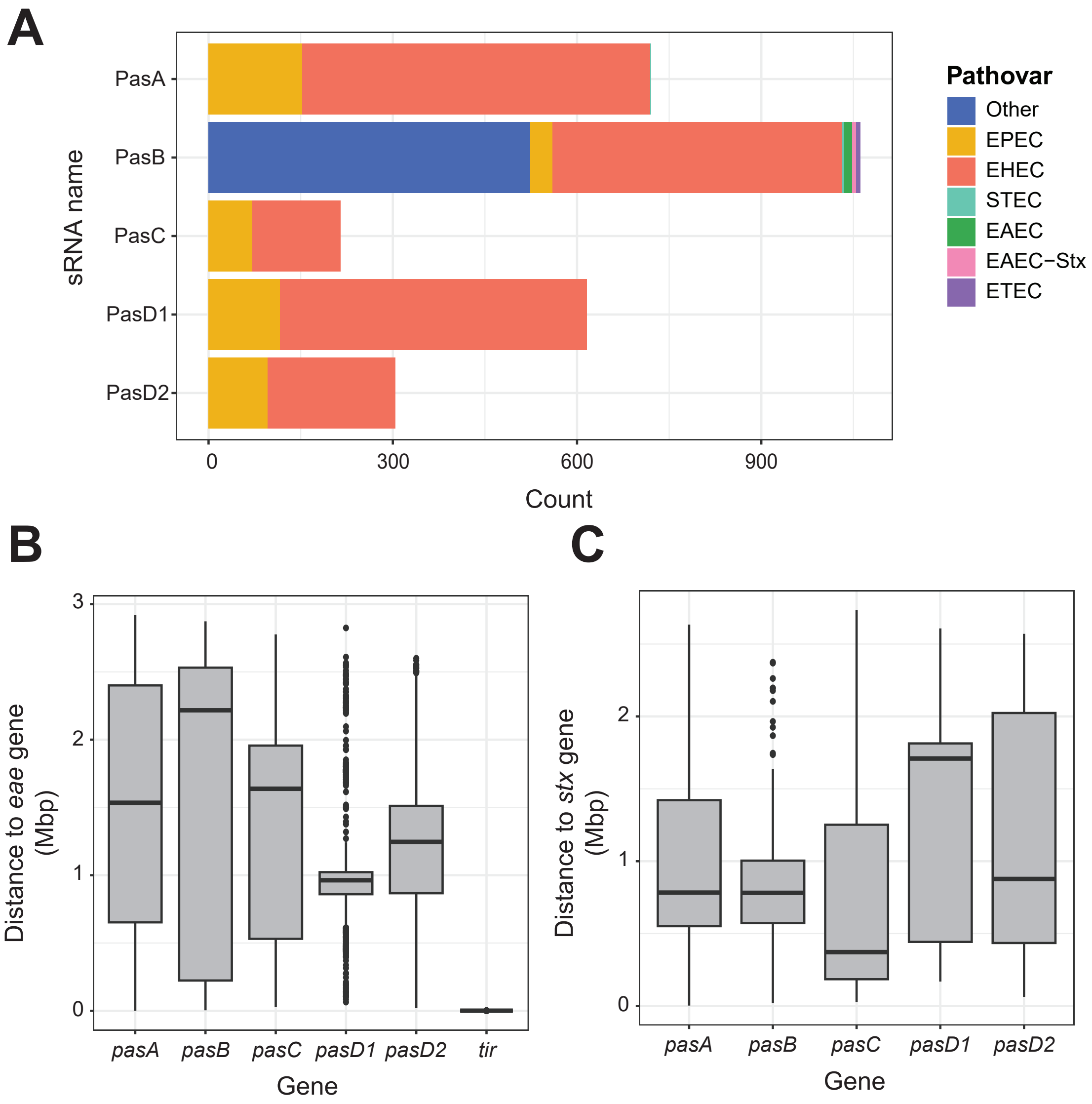


**Figure S3.** (A) Distribution of Pas sRNA genes across *E. coli* pathovars. *E. coli* isolates in the ‘pas_1362’ dataset were classified into one of 7 pathovars based on the presence of one or more definitive virulence factor genes: EPEC (*eae*), EHEC (*eae* and one of *stx1A*/*2A*), STEC (*stx1A/2A*), EAEC (*aggR*), EAEC-Stx (*aggR* and one of *stx1A/2A*), ETEC (*eltA/B*). Isolates classified as ‘Other’ lack any of the listed virulence factor genes but are not necessarily non-pathogenic. Boxplots showing the distances between Pas sRNA genes and EPEC and EHEC virulence factors (B) intimin, encoded by *eae*, and (C) Shiga toxins, encoded by *stx1A* or *stx2A*. As a control, the distance between *eae* and the translocated intimin receptor gene *tir*, which is encoded within the same pathogenicity island, is shown.

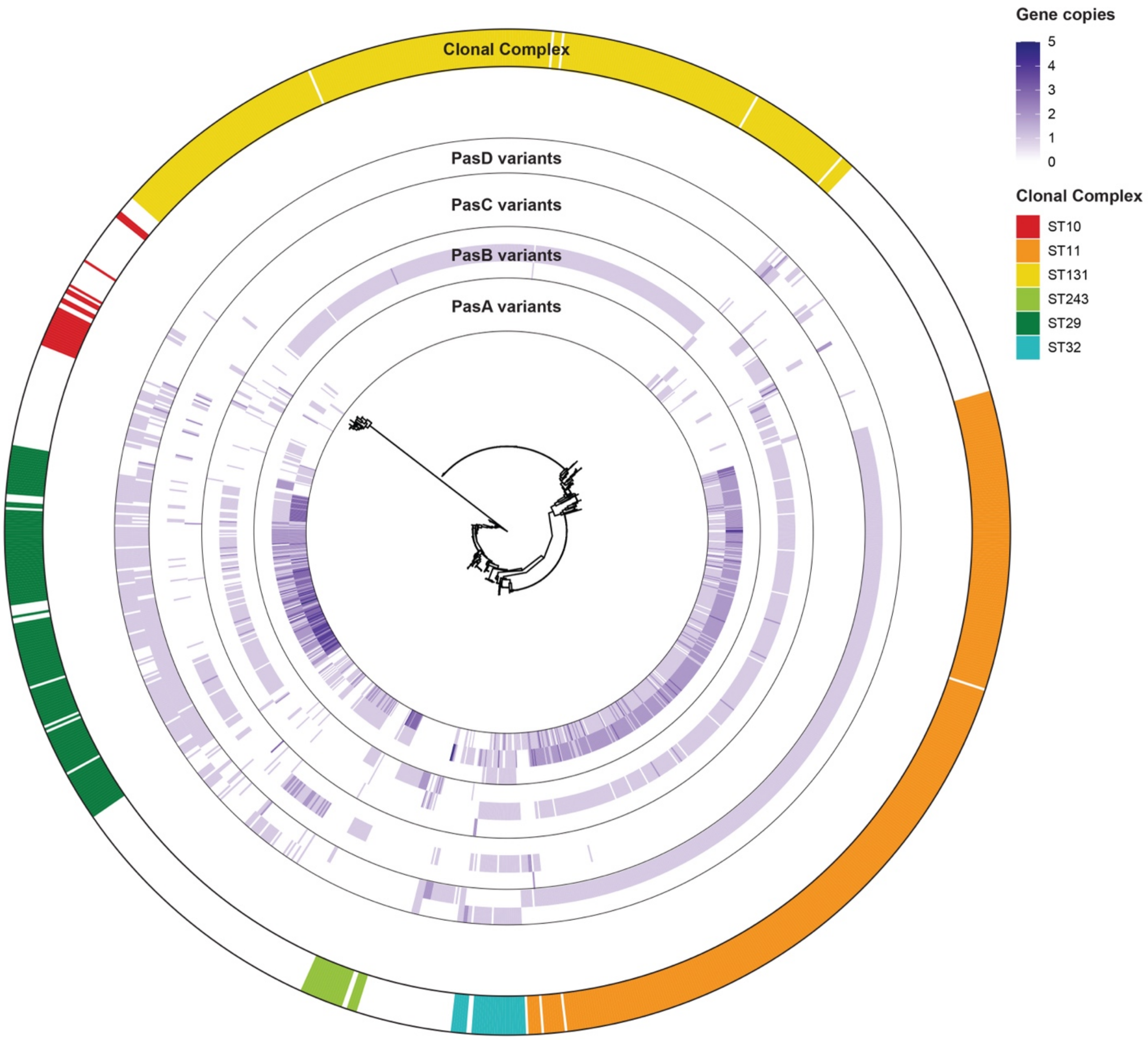


**Figure S4.** Maximum likelihood tree of 1,211 bacteria isolates from the 'pas_1362' dataset. Phylogenetic estimations were constructed from the nucleotide sequence of seven housekeeping genes from the Achtman multilocus sequence typing (MLST) scheme (*adk*, *fumC*, *gyrB*, *icd*, *mdh*, *purA*, *recA*). RAxML was used for maximum likelihood estimation using the 'GTRCAT' substitution model. The tree is rooted at the divergence of *Escherichia albertii* from *E. coli* isolates. The heatmap shows the gene copy number of Pas sRNA gene variants in each genome. The Pas gene variants are arranged (from inside to outside) in order of prevalence with the variant from EPEC strain E2348/69 listed first (e.g. PasA, PasA-2, PasA-3). The outer ring is colored based on the assigned MLST clonal complex of each isolate. Only the top 6 most prevalent clonal complexes are labeled.

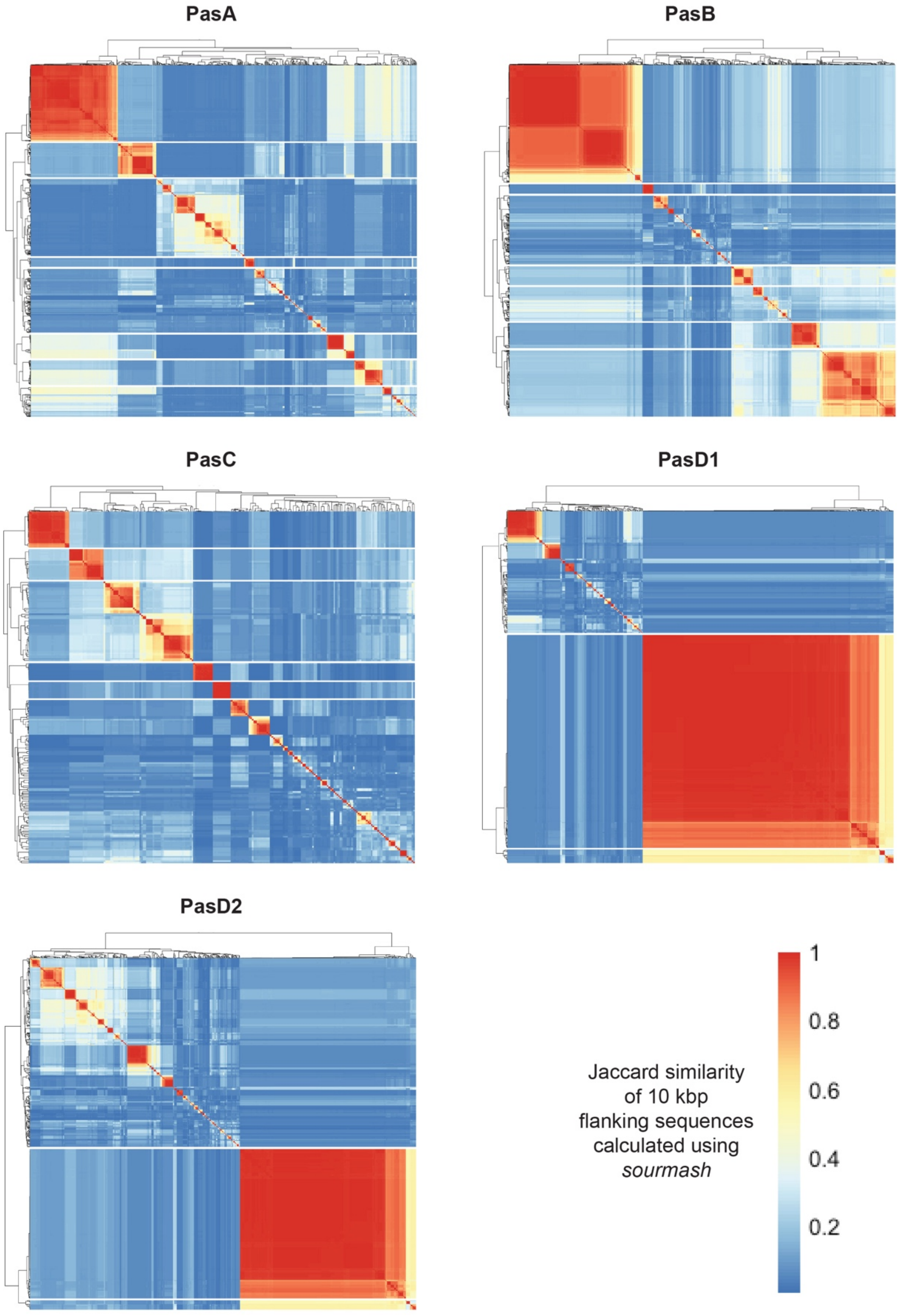


**Figure S5.** Clustering heatmaps showing the Jaccard similarity of the DNA sequence of 10 kbp flanking regions surrounding each Pas sRNA gene. Pairwise Jaccard similarity between each region was calculated using *sourmash* [4] to create a similarity matrix for all flanking regions for a given Pas sRNA. Briefly, the DNA sequence of the region is broken into 1,000x 31 bp kmers and the overlap in identical kmers between regions is calculated Agglomerative hierarchical clustering was performed using the similarity matrix to group regions based on the similarity of their nucleotide sequence. Cluster number was chosen so that the smallest cluster contained no less than 5% of the total regions in the dataset. Each row and column of the heatmap represents a single Pas sRNA flanking region. A representative locus from the most prevalent cluster for each Pas sRNA is depicted in **Fig. 4**.

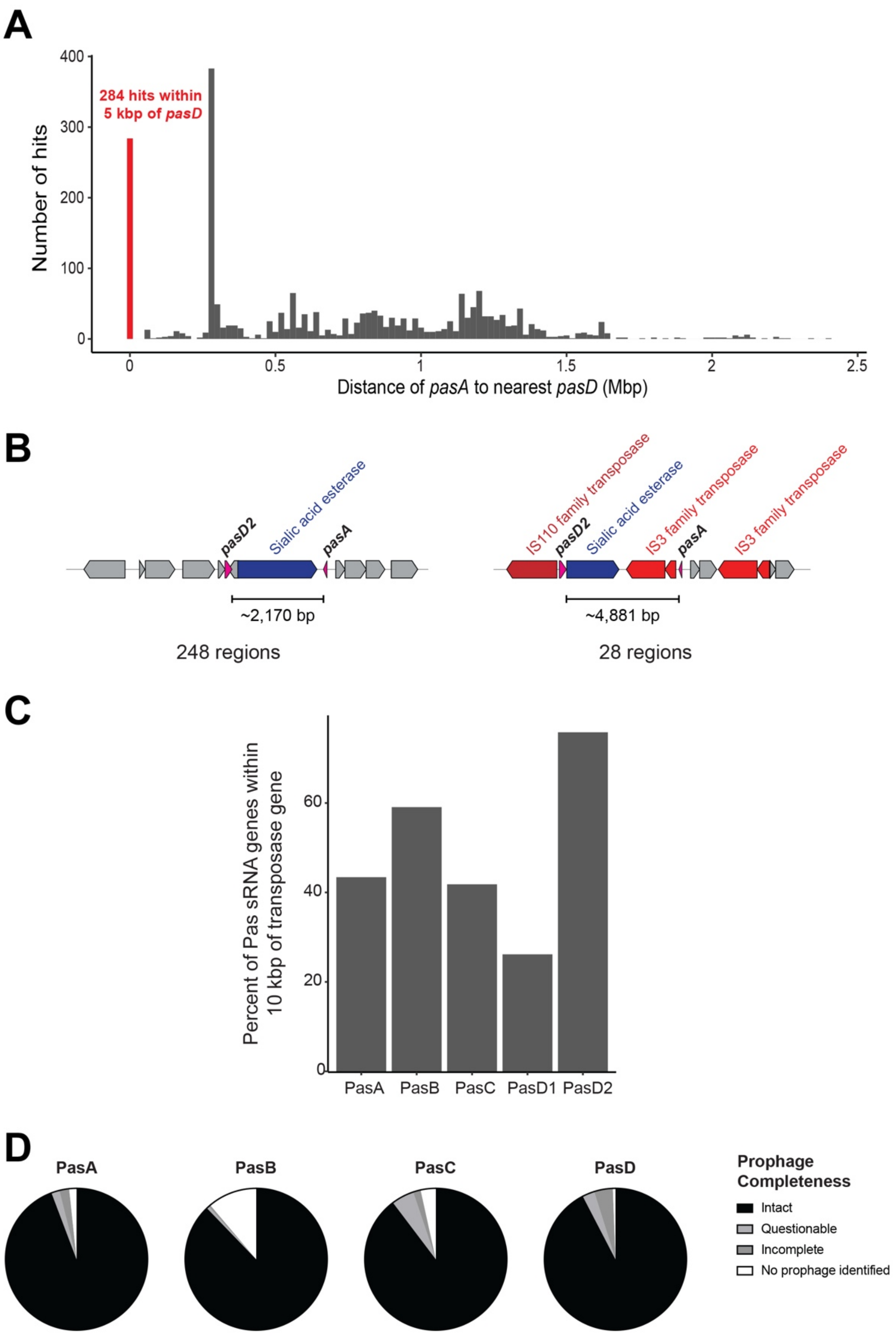


**Figure S6.** Positions of *pas* genes. (A) Histogram of the distance between each *pasA* hit in the 'pas_1362' dataset and the nearest *pasD* gene. Hits within 5000 bp (n = 284) are highlighted in red. (B) Gene arrangement found in the two most common synteny patterns for regions encoding both *pasA* and *pasD2*. The most common arrangement (left, n = 248) consists of *pasA* and *pasD2* flanking either side of a phage sialidase gene separated by ~2,170 bp. The second most common arrangement (right, n = 28) is similar to the former, except with several transposases interrupting the region and increasing the distance between *pasA* and *pasD2* to ~4,881 bp. (C) Bar graph of percentage of Pas genes within 10 kbp of a transposase gene. Transposase genes were identified based on the gene ontology annotation of coding sequences. (D) Pie charts show the percentage of *pas* sRNA genes from the ‘pas_1362’ dataset located within annotated prophage regions. Prophage regions were identified and classified based on completeness by PHASTER. Genes that did not map to an identified prophage region are labeled as ‘No prophage identified’.

**A**

| Gene | N Chromosomes Present | N Chromosomes Duplicate | Mean Distance (bp) | Min. Distance (bp) |
|---|---|---|---|---|
| *pasA* | 746 | 634 | 467,265 | 22,551 |
| *pasB* | 1,107 | 58 | 720,397 | 28,355 |
| *pasC* | 220 | 54 | 939,510 | 48,389 |
| *pasD* | 736 | 203 | 1,721,238 | 74,341 |

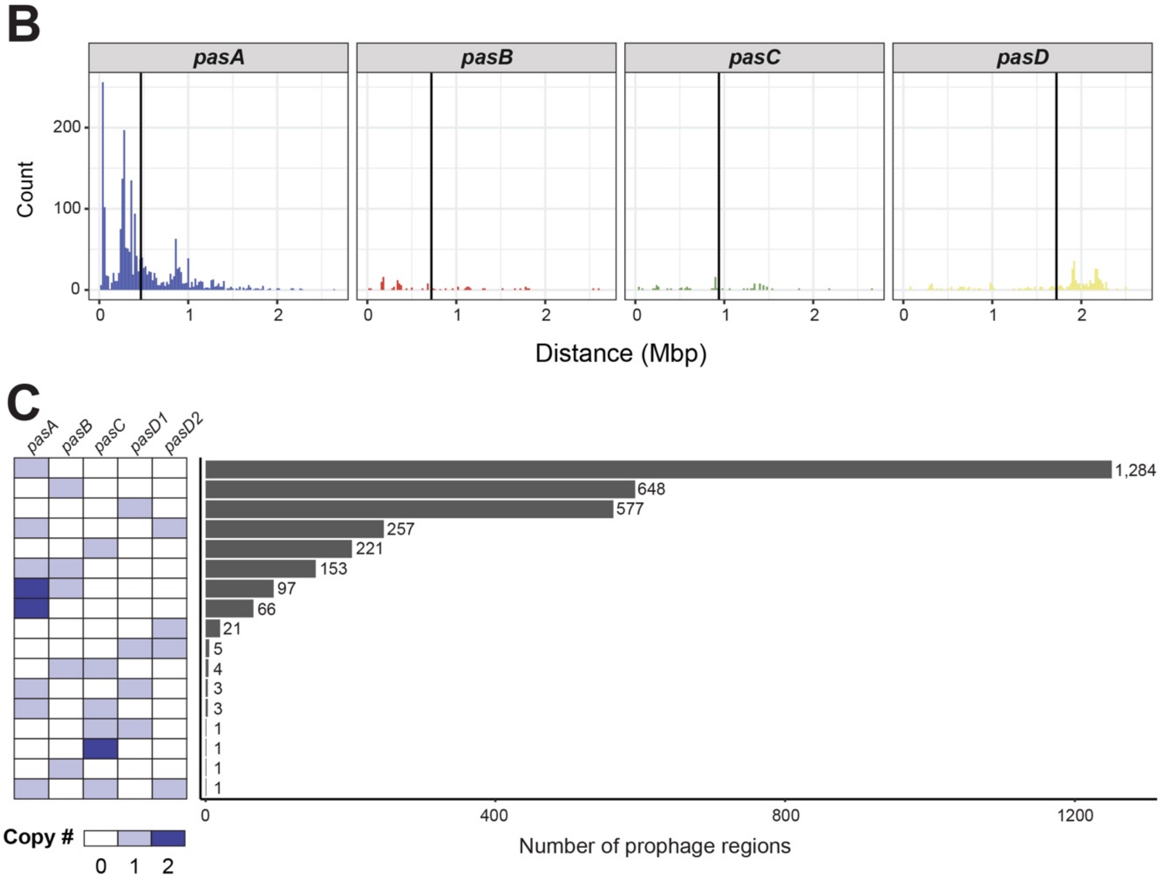


**Figure S7.** *pasA* is more frequently found in multiple copies compared to other Pas sRNA genes. (A) Table showing metrics characterizing the distance between Pas sRNA copies on the same chromosome. 'N Chromsomes Present' = total number of bacterial chromosomes in the 'pas_1362' dataset encoding each gene; 'N Chromosomes Duplicate' = number of chromosomes encoding more than one copy of a Pas gene; 'Mean Distance' = mean distance between copies of the same Pas sRNA gene when encoded on the same chromosome; 'Min. Distance' = the smallest distance between copies of the same Pas sRNA gene encoded on a chromosome. (B) Histograms showing the distances between duplicate copies of each Pas sRNA gene. Vertical black lines indicate the mean distance for each gene. (C) Pas sRNAs are typically found in singlicate within prophage regions on the bacterial chromosome. Bar graph showing the number of prophage regions containing different patterns of Pas sRNA genes indicated by the heatmap on the left.

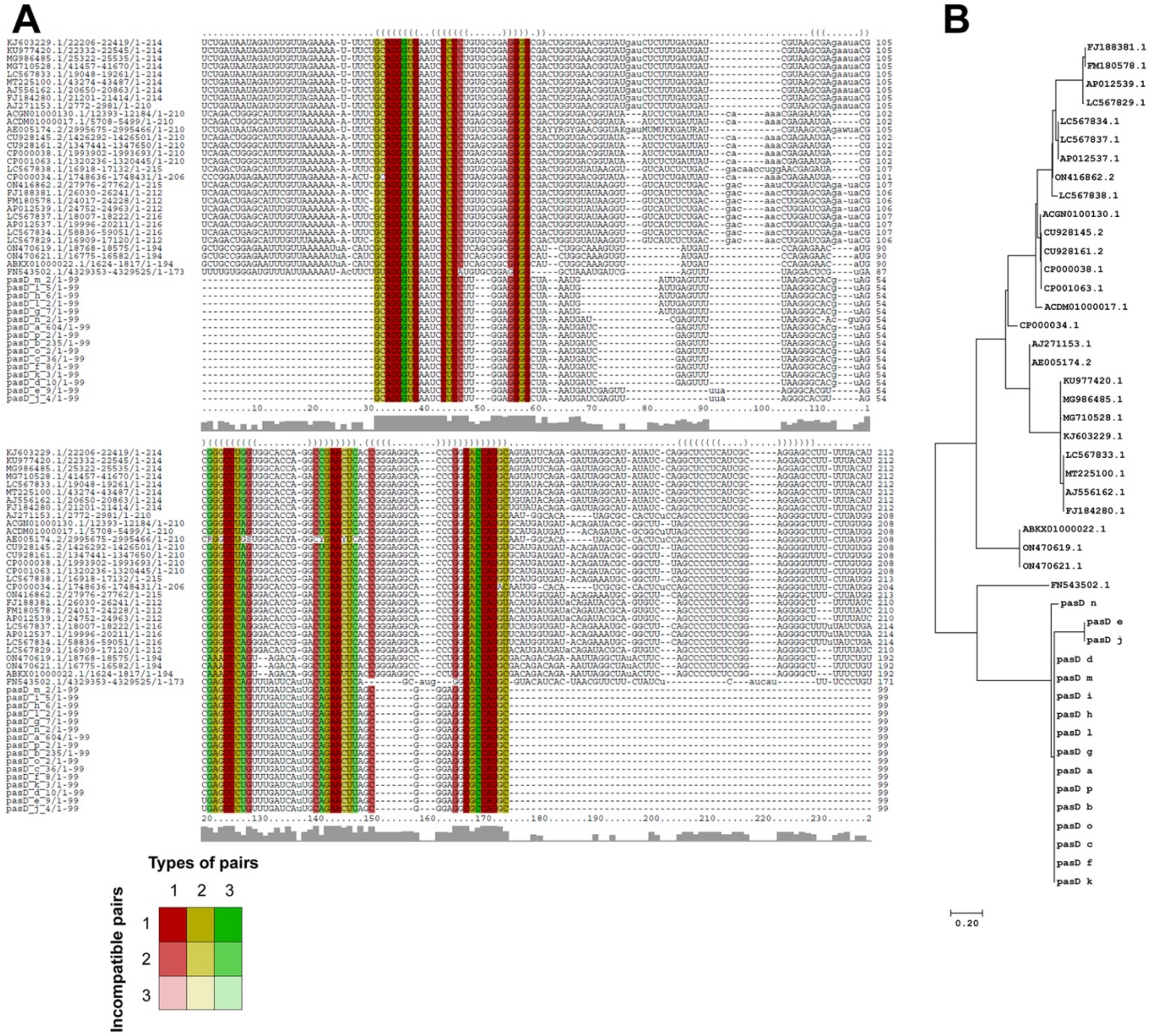


**Figure S8.** Similarity between PasD and STnc100 and corresponding evolutionary tree. Coloring scheme for alignment is as in (A). The evolutionary history was inferred by using the Maximum Likelihood method and Kimura 2-parameter model [2]. The tree with the highest log likelihood (-1198.69) is shown. Initial tree(s) for the heuristic search were obtained automatically by applying Neighbor-Join and BioNJ algorithms to a matrix of pairwise distances estimated using the Maximum Composite Likelihood (MCL) approach, and then selecting the topology with superior log likelihood value. A discrete Gamma distribution was used to model evolutionary rate differences among sites (5 categories (+G, parameter = 0.6782)). The tree is drawn to scale, with branch lengths measured in the number of substitutions per site. This analysis involved 46 nucleotide sequences. All positions with less than 50% site coverage were eliminated, i.e., fewer than 50% alignment gaps, missing data, and ambiguous bases were allowed at any position (partial deletion option). There were a total of 168 positions in the final dataset. Evolutionary analyses were conducted in MEGA11 [3].

### Supplementary tables

**Table S1.** (A) Filtered nucleotide BLAST search results. The query for the search was the DNA sequence for five Pas sRNA genes - pasA, pasB, pasC, pasD1, pasD2. The search was performed using the NCBI 'prok_complete_genomes' database accessed on April 18, 2025. The output format and column headers are standard from the BLAST+ suite '-outfmt 6'. (B) Table showing the Pas gene copy number (‘PasA’, ‘PasB’, ‘PasC’, ‘PasD1’, and ‘PasD2’), molecule length in base pairs (‘length’), and molecule type (‘mol.type’) of each nucleotide molecule represented in the ‘pas_1362’ dataset. ‘sseqid’ lists the NCBI accession number of each molecule.

**Table S2.** Linkage of Pas sRNA genes within bacterial genomes. (A) Pairwise Phi coefficients describing the presence or absence of Pas sRNAs within a given bacterial genome in the ‘pas_1362’ dataset. (B) Copy number of *pasA*, *pasB*, *pasC*, and *pasD* in each genome in the ‘pas_1362’ dataset. (C) Count table of the patterns of Pas sRNAs present in each *E. coli* genome, stratified by pathovar classification.

**Table S3.** Pas gene repertoires of notable *Escherichia* and *Shigella* strains.

**Table S4.** Percentages of Pas sRNA gene hits within 10 kbp of various genes of interest. Genes of interest were identified by searching ‘gene’ features in associated Genbank annotation files for specific keyword terms.

**Table S5.** Association of Pas sRNA genes with other sRNA and CSE hits predicted by *cmsearch* and *camscan*. The localization of other sRNAs and CSEs in proximity to Pas genes was analyzed in the flanking regions within 10 kbp of Pas genes in prophage and bacteriophage genomes. To compare RNA structured candidates with Rfam covariance models (CMs), and to perform structure-aware homology searches, we used *cmscan, cmsearch* (Infernal, version 1.1.5 [5]). By default, the inclusion threshold requires an E-value of 0.01 or less. Known sequence profiles and CMs were downloaded from the Rfam database [6]. The “tRNA_dist” worksheet presents the distributions of distances between Pas sRNA genes and tRNA hits for PasA and PasC, where such hits occur most frequently.

**Table S6.** (A) Summary of 43 unique bacteriophages with at least one Pas sRNA gene detected by both BLAST and cmscan with cutoffs of e<0.005 and qcov>75%. (B) Results of blastn search of Pas sRNA genes in the Millard Lab Bacteriophage database (May 03, 2025 version) filtered for results with query coverage >=75%. (C) Results of blastn search of Stx1A and Stx2A genes in the Millard Lab Bacteriophage database (May 03, 2025 version).

### Supplementary References


1. Pearl Mizrahi, S., Elbaz, N., Argaman, L., Altuvia, Y., Katsowich, N., Socol, Y., Bar, A., Rosenshine, I. and Margalit, H. (2021) The impact of Hfq-mediated sRNA-mRNA interactome on the virulence of enteropathogenic *Escherichia coli*. *Sci. Adv.*, **7**, eabi8228.
2. Kimura, M. (1980) A simple method for estimating evolutionary rate of base substitutions through comparative studies of nucleotide sequences. *J. Mol. Evol.*, **16**, 111-120.
3. Tamura, K., Stecher, G. and Kumar, S. (2021) MEGA11: Molecular Evolutionary Genetics Analysis Version 11. *Mol. Biol. Evol.*, **38**, 3022-3027.
4. Pierce, N.T., Irber, L., Reiter, T., Brooks, P. and Brown, C.T. (2019) Large-scale sequence comparisons with sourmash. *F1000Res*, **8**, 1006.
5. Nawrocki, E. and Eddy, S.R. (2013) Infernal 1.1: 100-fold faster RNA homology searches. *Bioinformatics*, **29**, 2933-2935.
6. Ontiveros-Palacios, N., Cooke, E., Nawrocki, E.P., Triebel, S., Marz, M., Rivas, E., Griffiths-Jones, S., Petrov, A.I., Bateman, A. and Sweeney, B. (2025) Rfam 15: RNA families database in 2025. *Nucleic Acids Res.*, **53**, D258-D267.
